# Targeted Nanoparticles: an Innovative Modality in the Treatment of Cancer

**DOI:** 10.64898/2025.12.01.691687

**Authors:** Hyung-Gyoo Kang, Bryon Upton, Ryan W. Holly, Racheal Upton, Sheetal Mitra, Jean-Hugues Parmentier, Ann F. Mohrbacher, Yong-Mi Kim, Timothy J. Triche, Jon O. Nagy

## Abstract

Despite progress made in the development of anticancer therapeutics, traditional small-molecule chemotherapeutics often struggle to overcome toxicity, efficacy, and off-target effects. These specific issues can be overcome by either encapsulating the drug or by targeting it directly to the tumor cell. Here, we describe a novel targeted nanoparticle (referred to as a nano-antibody-drug conjugate Targeted Nanosphere or nADC/TNS), based cancer therapeutic platform that can improve the efficacy of a broad range of existing therapeutics. Targeting is antibody-directed, as with antibody-drug conjugates (ADCs). Still, the payload per antibody is vastly greater by orders of magnitude (a thousand for nADC/TNS versus two to eight for ADCs). The nADC/TNS consists of an approximately 80 nm drug-filled nanoparticle composed of phospholipids, cholesterol, and UV cross-linkable diacetylene lipids. We describe the preparation, characterization, and evaluation of nADC/TNS as a novel, versatile, and effective treatment modality for cancer and potentially other diseases. This report focuses on data with nADC/TNS variants NV101 (anti-CD99 targeted, doxorubicin-filled), NV102 (anti-CD19 targeted, doxorubicin-filled), and NV103 (anti-CD99 targeted, irinotecan-filled). We investigated NV101 and NV103 in a mouse model with implanted and metastatic Ewing tumors (ES). NV101 demonstrated significant tumor burden reduction while NV103 induced complete ablation of ES tumors. NV102 demonstrated complete ablation of chemotherapy-resistant relapsed adult lymphocytic leukemia (ALL). These results document the potential superior efficacy of antibody-targeted nanoparticles containing a variety of small-molecule payloads, compared to their free molecule equivalents.

## INTRODUCTION

The development of rationally designed nanoparticle drugs to treat challenging diseases began in the 1960s [1]. Since their inception, nanoparticle-based drugs have advanced dramatically, improving drug delivery while reducing treatment-related toxicity. Examples include doxorubicin HCI liposomal injection (Doxil^TM^), paclitaxel protein-bound particles for injectable suspension (Abraxane^TM^), paclitaxel liposome for injection (Lipusu^TM^), irinotecan liposome injection (Onivyde^TM^), cytarabine and daunorubicin liposome for injection (Vyxeos^TM^), and liposomal cytarabine (DepoCyt^TM^), with several new candidates currently in clinical trials [2]. Nanoparticle formulations include liposomes, dendrimers, polymer and co-polymer conjugates. Their therapeutic cargo may be either passively (untargeted) or actively targeted to tumor cells. Notwithstanding, to date, all the clinically approved nanoparticle drug carriers are *untargeted* particles. In contrast, antibody-drug conjugates (ADCs) leverage the specificity of antibodies to deliver potent cytotoxic agents directly to tumor cells. However, ADCs are limited to a handful of drug molecules delivered per antibody molecule internalized by the tumor cell.

An antibody-targeted nanoparticle containing thousands of small molecules per nanoparticle offers the theoretical possibility of preferentially delivering more drug to tumor cells while bypassing normal tissues that do not express the target antigen, which should result in little or no treatment-related toxicity.

The targeted nanoparticle (TNP) described here, is in essence is a *novel format* of nano-antibody-drug conjugate Targeted Nanosphere (nADC/TNS). The basic structure is an antibody-targeted particle that is a ***h****ybrid **p**olymerized **l**iposomal **n**anoparticle* (HPLN) composed of conventional phospholipids, pegylated-phospholipids, cholesterol, and polydiacetylene “PDA” photo-cross-linked polymers. While crosslinked PDA polymers have been incorporated into potential pharmaceutical products (see below), none are in routine clinical use in humans. The advantage of introducing polymer lipids into a liposome formulation is that it imparts rigidity to the nanoparticle, preventing fusion with the cell membrane, unlike many conventional liposomal formulations. It is known that membrane fusion stimulates MDR1 and similar drug efflux effects [3]. In contrast, nADC/TNSs are exclusively incorporated into the cell via endocytosis and lysosomal fusion, bypassing such mechanisms, thereby enhancing direct drug delivery after lysosomal-mediated release from the nanoparticle.

While advances in ‘precision oncology’, with tumor-targeted therapies like immunotherapeutics (e.g., pembrolizumab (Keytruda^TM^), nivolumab (Opdivo^TM^), and ipilimumab (Yervoy^TM^) and kinase inhibitors (e.g., MET, RAS, EGFR) designed to treat specific tumors and their particular genetic defect (e.g., trastuzumab (Herceptin^TM^) and HER2 amplified breast cancer), patients whose tumor lacks these specific targets still receive conventional small molecule therapy, usually as a free drug, which often results in *dose-limiting toxicity* (DLT) that prevents use of *maximally effective doses* (MED). There is thus an urgent medical need to develop more effective therapy for cancers that do not benefit from targeted therapies, which unfortunately is true of the majority who eventually relapse after initial favorable clinical responses. Efforts to find new effective drugs with fewer side effects that can lead to complete clinical remission without significant toxicity or adverse events are one of the most, if not the most, important goals of cancer therapy development.

At the same time, revisiting existing, effective drugs that ordinarily exhibit unacceptable toxicity when delivered as a free drug may yield a better understanding and deeper insight into the specific pathway and biological targets they disrupt, and may lead to their toxicity. Understanding the mechanisms of toxicity can further guide the development of innovative drug delivery platforms, including nADC/TNS, designed to specific targets while minimizing exposure to normal or healthy tissues, reducing side effects, and improving safety.

Here we do so to demonstrate that such drugs, when preferentially delivered to the tumor, can minimize or eliminate DLT while significantly increasing the *maximum tolerated dose* (MTD) [4,5]. In this study, we show that *tumor-specific targeting* combined with drug-loaded nanoparticles can achieve this goal. nADC/TNSs are a novel nanoparticle formulation (also referred to as a novel format conjugate) that enables this tumor-specific targeting of a payload consisting of thousands of small molecules of choice, resulting in dramatic tumor response with no discernible toxicity.

As a novel technology, utilizing tumor-specific targeting agents on the surface of drug-encapsulated nanoparticles can: (a) ***preferentially*** deliver the drug to cancer cells while sparing normal cells and tissues, and (b) deliver lower ***systemic*** cytotoxic doses to minimize both initial and long-term morbidity.

Improving on the high tumor cell specificities already seen for ADCs, an nADC/TNS can provide a significantly greater on-target drug-to-antibody ratio (DAR) than an ADC, namely, a *thousand* drug molecules per nanoparticle-conjugated antibody versus only 2 to 8 conjugated drug molecules per ADC antibody [6]. Increasing the amount of drug delivered *per targeting antibody* will improve the overall efficacy and reduce any toxic side effects coming from the antibody alone. The desire to reduce dose-limiting toxicity of ADCs is moving the field now to investigate new high drug-content loadable scaffolds that are conjugated to single antibodies. The nADC/TNS technology is able to move this strategy to an even higher level.

In addition, the polyvalency of having a number of antibodies attached to a single nanoparticle enhances the overall avidity of nanoparticles to target tumor cells, compared to a single ADC molecule. Moreover, in a hollow-shell nanoparticle, the encapsulated drugs are not chemically conjugated to the carrier, as they are in the ADC assemblies. Once released from the carrier, no *in situ* chemical decoupling is required for the drug to become fully bioavailable.

The need for sophisticated drug delivery systems has facilitated the emergence of polymeric nanoparticles at the forefront of promising new technologies [7]. The potential pharmaceutical uses for PDA-containing liposomes, micelles, nanofibers, or other nanovesicles include vaccine applications [8], cancer drug delivery [9–14], oligonucleotide delivery [10, 15–18], and a variety of other potential disease treatments [19–23]. While these applications were tested in cell-based assays, numerous PDA-containing liposomes and coatings for nanoparticles have been tested in mice [24–34], Zebrafish [7], and rats [35, 36] with no adverse effects. These studies demonstrate the utility, biocompatibility, and relative safety in living organisms of the polydiacetylene polymers [37].

This report describes nADC/TNP preparation, their characterization, and evaluation in pre-clinical models. For the initial assessment of the nADC/TNP, we chose adult leukemia (ALL) and Ewing Sarcoma (ES) cells to validate the core technology *in vitro* and tumor burden reduction models in transgenic mice. ALL and ES were chosen because they generally lack dense tumor-associated stroma that can block access to tumor cells, and because they are well vascularized, ensuring ready access of systemically administered nADC/TNP to tumor cells [38]. These tumor types are thus ideal models to evaluate the efficacy of nADC/TNS-based therapies. The chemotherapeutics encapsulated were doxorubicin (Dox) or irinotecan (Ir), and the targeting antibody was either *anti*-CD99 (NV101, NV103) or *anti*-CD19 (NV102).

In this paper, we present the results from two *in vivo* studies in a *NOD/SCID* transgenic mouse model, in which we evaluated NV101 and NV103 for efficacy in reducing tumor burden and survival against ES. In addition, we evaluated NV102 for its effectiveness in ablating treatment-resistant adult leukemia.

## RESULTS

### Hybrid Polymerized Liposomal Nanoparticles (HPLN)

The unique nanoscale particle shell, referred to as the HPLN, is the base structure of the targeted drug assembly that will become the nADC/TNS and is illustrated in **Figure 1**, showing a schematic diagram (a), a negative-stained TEM image (b), and a cryo-EM image (c) of an individual particle. This type of liposome is a new class sometimes referred to as nanosomes or polymerosomes [39].

**Figure 1.**
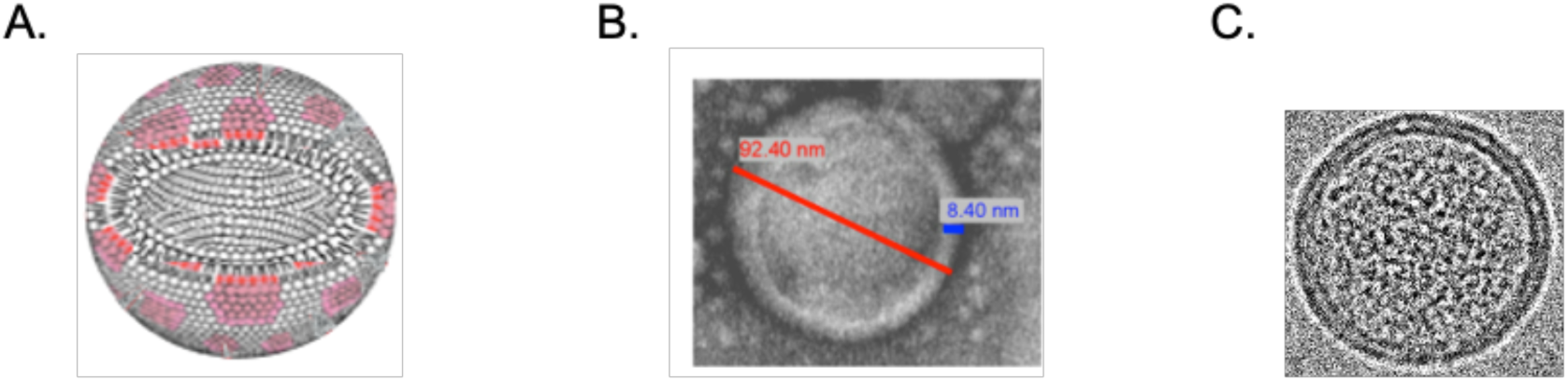
**A.** A schematic of the HPLN showing the distinct regions of polymerized lipid patches (pink) surrounded by ordinary phospholipids (white); **B.** a negative stain electron microscopic image of a single HPLN with a cross-sectional bar to estimate the dimensions; and **C.** an HPLN cryo-EM image.

The components of the HPLN liposome precursor to the nADC/TNP are depicted in **Figure 2**.

**Figure 2.**
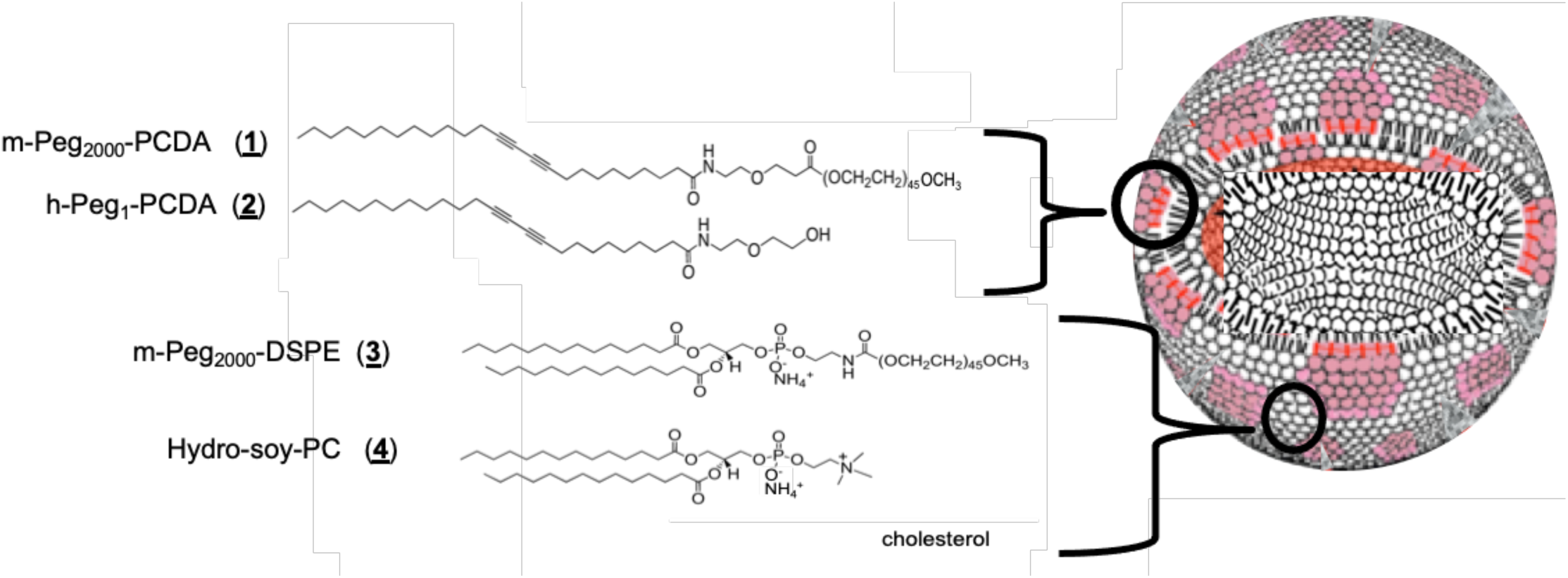
The lipid components and cholesterol that comprise the HPLN shell. The PDCA lipids segregate into patches or islands that allow the polymer, polydiacetylene, to form after exposure to UV light.

For the current study, to produce the final nADC/TNS, NV101, NV102, and NV103, the HPLN/drug formulations were coated with monoclonal human antibodies toward leukemia cells (*anti*-CD19, NV102) and ES cells (*anti*-CD99 NV101 and NV103). The general nADC/TNS construction strategy scheme is depicted in **Figure 3**.

**Figure 3.**
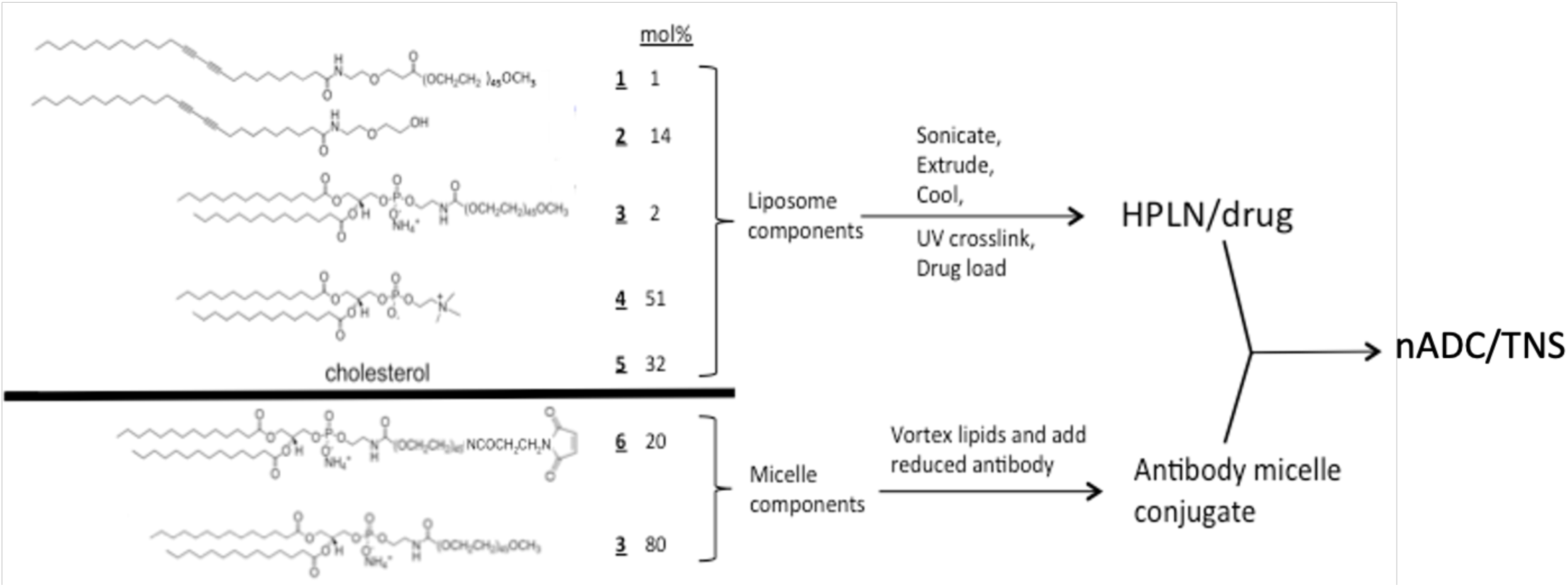
Schematic of the nADC/TNS preparation process. The drug-loaded HPLNs are prepared as a storable solution. In a separate process, the antibody-labeled micelles are constructed and admixed to the loaded HPLNs to complete the nADC/TNP assembly.

To prepare an nADC/TNP particle, we started with a standard stealth liposomal formulation, such as found in Doxil^TM^. The Doxil^TM^ liposome formulation consists of hydrogenated soy PC *<u>4</u>* (where the primary component is distearoylphosphatidylcholine (DSPC)), cholesterol, and polyethylene glycol-distearoylphosphatidyl ethanolamine *<u>3</u>* (m-PEG2000-DSPE) in a molar proportion of 57.5: 37.5: 5. To this formulation, we added varying amounts of the simple derivatives of the unsaturated and polymerizable lipid 10,12-pentacosydiyanoic acid (PCDA) [40].

We chose a very short head-group mono-ethylene glycol amide derivative, h-PEG_1_-PCDA ***<u>2</u>***, because it is readily crosslinked by UV light exposure. The other desirable physical properties include: a) good aqueous dispersion properties when mixed with charged lipids; b) being uncharged, it will not alter the overall surface charge or Zeta potential, and c) the small polar head will not sterically interfere with the recognition of surface-attached targeting agents, co-displayed on the nADC/TNP surface.

After evaporating the lipid mix to a thin film, bath sonication while heating the waxy solid results in crude liposomes in a broad size range. Repeated passage of the liposomes over an 80 nm extrusion membrane, also while heating, gives liposomes of a tight, homogeneous size population (ca. 79-81 nm). Upon cooling, the liposomes do not change their overall size, but the lipids phase separate on the surface of individual particles into sections of similar hydrocarbon chain length. Because the *polymerizable* diacetylene lipids are longer in chain length (C-25) than the other PC lipids (C-16 and 18), lipid islands or patches in the liposome membrane occur [41], not unlike the lipid rafts that occur normally in cell membranes. Now clustered together in close proximity, these diacetylene rafts readily polymerize after brief UV irradiation into polydiacetylene patches [42], leading to deeply colored particles (see pink polymer patches depicted in **Figures 1** and **2**).

The h-PEG_1_-PCDA (***<u>2</u>***) is the lipid responsible for the resulting polydiacetylene polymer. While this lipid alone will readily form liposomes, it is uncharged, and the h-PEG_1_-PCDA particles are not colloidally stable and will quickly drop out of solution. When mixed with other lipids such as zwitterionically charged PC (*<u>4</u>*) and polyethylene glycol (PEG) terminated PC lipids (*<u>3</u>*), the resulting liposomes are somewhat more colloidally stable. By varying the amount of h-PEG_1_-PCDA (*<u>2</u>*), we found that at 14 mol% concentration relative to the other components, these liposomes gave a robust amount of polymer, as evidenced by a deeply colored liposome solution after cooling and brief UV irradiation [43]. Increasing the molar percentage above 14 mol% seems to lead to slight increases in overall average particle diameters (**Table 1**).

**Table 1.** A comparison of HPLN formulations that vary the relative amounts of h-Peg_1_-PCDA lipid from 14 to 44%, formulated by reducing the proportional quantity of hydro-soy PC lipid.

| Percent polymer in HPLN<br>(determined by the relative<br>mole ratio of h-PEG<sub>1</sub>-PCDA<br>lipid to the other liposome<br>components). | Absorbance<br>at 640 nm | Average<br>particle size | Amount of<br>doxorubicin<br>loaded (mg<br>Dox/mg HPLN) |
| --- | --- | --- | --- |
| 14% | 0.716 | 97nm | 0.13 |
| 24% | 1.335 | 100nm | 0.07 |
| 34% | 1.828 | 114nm | 0.06 |
| 44% | 1.975 | 126nm | 0.04 |

In the initial formulations of polymerizable liposomes containing lipids *<u>2</u>, <u>3</u>, <u>4</u>*, and cholesterol, once the lipids segregated into areas of similar chain length populations, particle aggregation became evident upon several weeks of storage. We hypothesized that upon lipid phase segregation into the ‘islands’, the h-PEG_1_-PCDA lipid polymer patches could act as adhesive sites and promote inter-particle adherence, leading to colloidal instability and ultimately precipitation. This aggregation was remedied by inclusion of a small amount (1 mol%) of the crosslinkable *pegylated* m-PEG_2000_-PCDA molecule (***<u>1</u>***). This lipid, in principle, should diffuse into the h-PEG_1_-PCDA islands since it is of similar hydrocarbon tail length and provides a steric barrier, inhibiting any particle-particle aggregation promoted by the h-PEG_1_-PCDA patches alone. The final HPLN formulation is depicted in **Figure 3**. Particles containing pegylated m-PEG_2000_-PCDA (***<u>1</u>***) are stable at 4°C for months, with little observed precipitation.

The HPLNs can also be made to fluoresce intrinsically. Upon initial polymerization, the “blue” form of the polydiacetylene polymer is non-fluorescent, but after heat treatment, the blue vesicles change to red-pink color, resulting from the formation of a fluorescent fluorophore [44]. The color change is a result of a subtle conformational change in the polymer backbone, leading to the fluorescent polymer form. The fluorescence emission of the HPLNs gives an emission spectrum that is centered at 635 nm with a broad and complex excitation spectrum from 480 to 580 nm, enabling simple detection by FACS and other fluorescence-based assays.

### Drug loading into HPLNs

When the initial HPLN-lipid mixture is sonicated in ammonium sulfate, the liposomes can be actively loaded with amine-containing drugs such as doxorubicin or irinotecan. This is possible due to the pH gradient that is established across the particle lipid membrane, once the exterior ammonium sulfate is removed by dialysis [45]. In terms of drug loading, we have found that even in the polymerized form, the nanoparticles can be actively drug-loaded with doxorubicin or irinotecan. Loading the drug post-polymerization into HPLNs is crucial because it prevents exposing the therapeutic agent to potentially damaging UV light. In earlier studies, we found that the concentration of polymerizable PCDA lipids required for drug loading to occur, required a polymer content of approximately 15 mol percent (or less) [46]. The non-encapsulated drug is removed by continuous TFF dialysis filtration until the effluent is free of drug checked by spectrophotometric analysis.

As seen in **Table 1**, as the relative percentage of crosslinkable lipid h-Peg1-PCDA is increased, the amount of doxorubicin that can be loaded drops off as expressed as a ratio of drug to particle concentration. Like the process described for doxorubicin encapsulation, irinotecan can also be actively loaded into ammonium sulfate-loaded HPLNs.

**Table 2** shows the initial liposome size (after extrusion), upon cooling and UV crosslinking, and finally upon doxorubicin or irinotecan loading.

**Table 2.** Average particle size and PDI throughout initial liposome formation, polymerization, and drug loading. **\***PDI is a measure of the size homogeneity of a particle population in a sample. If a particle sample is becoming aggregated, the PDI gets large and approaches a value of **1.0**. PDI values in the range of 0.3 or greater indicate a **heterogeneous** size population in the sample. Values between 0.2 and 0.1 are considered very good, and those of 0.1 or less are excellent, suggesting a homogenous population with a narrow size distribution.

| Particle preparation step | Process description | Size: average diameter/polydispersity (PDI)* |
| --- | --- | --- |
| 1 | Initial liposome prep (pre-polymerized) | 79 nm/0.10 |
| 2 | Polymerization to HPLN (prior to drug load) | 79 nm/0.12 |
| 3 | Load HPLN with Dox or Ir | 82 nm/0.13 |

#### HPLN/Dox and HPLN/Ir

Since doxorubicin is still a mainstay of the current treatment of leukemia and many other types of cancers [47], it was chosen as our initial nADC/TNS particle drug payload. As reported [46], the IC_50_ concentrations of *untargeted* Dox-loaded HPLNs, in three independent osteosarcoma tumor-derived cell lines, were at least 6-fold *lower* (more cytotoxic) than the conventional liposomal doxorubicin formulation. This boost in potency appears to depend on the PDA polymer content, since it is progressively lost as the h-PEG_1_-PCDA lipid component concentration is titrated down from an optimum of 15–20 mole percent [46].

The mechanism for this enhanced efficacy is not fully understood, but it may be a function of the superior drug release characteristics of the nADC/TNS nanoparticle, once inside the cell. While both drug carrier vehicles (HPLN/Dox and conventional liposome/Dox) showed approximately the same degree of *passive* drug release in neutral pH buffer, upon lowering the pH from 7.4 to 4.5, an increased drug release in the HPLN occurred by a factor of 1.5 (P =0. 01; 95% C.I. 1.1- to 2.1-fold) with evidence that this effect was enhanced at 37°C [46]. This enhanced drug release under acidic pH from encapsulated polymerized PDA liposomes was also observed in vesicles containing ampicillin [49] and amino-naphthalene [50].

Numerous reports demonstrated that partial polymerization of nanoparticles provide an enhanced stability of the encapsulated drug against passive leakage [48–51], when compared to the conventional liposome forms. This result suggests that at neutral pH, at physiological temperature, greater concentrations of drug would be expected to be delivered to target cells in the HPLN form, compared to the conventional liposome form.

The encapsulation amount of doxorubicin inside HPLN/Dox was measured to be 0.14 mg Dox/mg HPLN particle mass. This compares very closely to the commercial doxorubicin-encapsulating liposome Doxil^TM^ at a 0.13 mg Dox/mg liposome mass [52]. The irinotecan-loading into the HPLN was routinely determined to be between 0.1 and 0.15 mg Ir/mg HPLN particle mass. Using a NanoSight device to assess the actual number of HPLN particles in a given volume of solution and measuring the total drug concentration in this volume after particle rupture, we estimate the number of drug molecules per HPLN to be about 10,000-20,000.

### Construction of novel format conjugates (nADC/TNS)

The final step in the preparation of the nADC/TNS requires the appending of tumor-targeting antibodies. In our first generation of nADC/TNS (with encapsulated doxorubicin), we reported the utilization of a recombinant *anti-activated leukocyte adhesion molecule* (ALCAM, CD166) antibody for targeting [46]. This resulted in a *12-fold increase* in tumor cell cytotoxicity over the conventional Doxil^TM^ formulation.

Others have reported the targeting of lymphoma cells by *rituximab (Rituxan^TM^)* Fab-conjugated, polydiacetylene liposomes [24]. In this report, the targeted polymerized liposomes were significantly more effective than either the free drug or non-targeting liposome in inhibiting primary tumor growth and prolonging the graft survival. Both the *anti*-ALCAM targeted nADC/TNS and the *rituximab* targeted Fab liposomes were more effective in cell killing *in vitro* than the non-targeted forms.

For later generations of *nADC/TNS* our new methodology is described in **Figure 3**. The improvement allowed drug loading at a higher per-particle amount than described in our earlier report [46].

For the current study, to produce the final nADC/TNSs, NV101, NV102, and NV103, the HPLN/drug formulations were targeted with monoclonal human antibodies toward leukemia cells (*anti*-CD19) and ES cells (*anti*-CD99), respectively. The general nADC/TNS construction strategy scheme is depicted in **Figure 3**.

To obtain antibody-targeted nADC/TNS formulations, targeting antibodies were partially reduced to liberate sulfhydryl moieties from the cysteines in the protein structure [53]. The sulfhydryl groups were reacted with maleimide-terminated, pegylated lipids in pre-made micelles [54], **Figure 3**. By mixing two pegylated DSPE lipids (*<u>3</u>* and *<u>6</u>*) in aqueous buffer, a clear solution of micelle particles was readily formed. Adding the partially reduced antibodies to the micelle solution allowed a Michael-type addition reaction between the thiol of the antibody-cysteine and the maleimide moiety in the micelle, resulting in covalent coupling of the antibody to the micelle.

As a negative control, micelles were prepared by reacting cysteine with the maleimide lipids micelles. The antibody-labeled micelle was incubated with the drug-loaded HPLNs, and an efficient insertion of all the micelle lipids [54] into the HPLN membrane occurred, resulting in the antibody-labeled nADC/TNS.

### Choice of targeting antibody

Nanoparticle anticancer targeting strategies require the identification of a biomarker expressed on the surface of the tumor cell. Tumor-associated molecules expressed at higher levels than in normal tissues are sought since nanoparticles coated with antibodies recognizing these markers can, in principle, preferentially bind to tumor cells.

Early work [46] has shown that, unlike liposomal nanoparticles, nADC/TNPs are primarily taken up by cells via clathrin and/or caveoli-mediated endocytosis that is distinct from the uptake pathway for small molecule drugs, which are taken up by passive diffusion across cell membranes [55]. Thus, it is generally recognized that nanoparticles that enter the cell by endocytosis can enhance the efficacy of chemotherapeutic agents by circumventing the MDR pump, which otherwise significantly reduces the therapeutic activity of small molecule drugs. In addition, nADC/TNS intracellular uptake is dramatically enhanced by the nanoparticle surface-displayed antibodies, which provide active targeting to the tumor antigen. As seen in **Figure 4**, after 24 hours, massive cellular uptake of the nADC/TNS into discrete endophagocytic vesicles, at least some of which have fused with lysosomes is visible. It is clear from the image that the nanoparticles enter cells via the endocytotic pathway.

**Figure 4.**
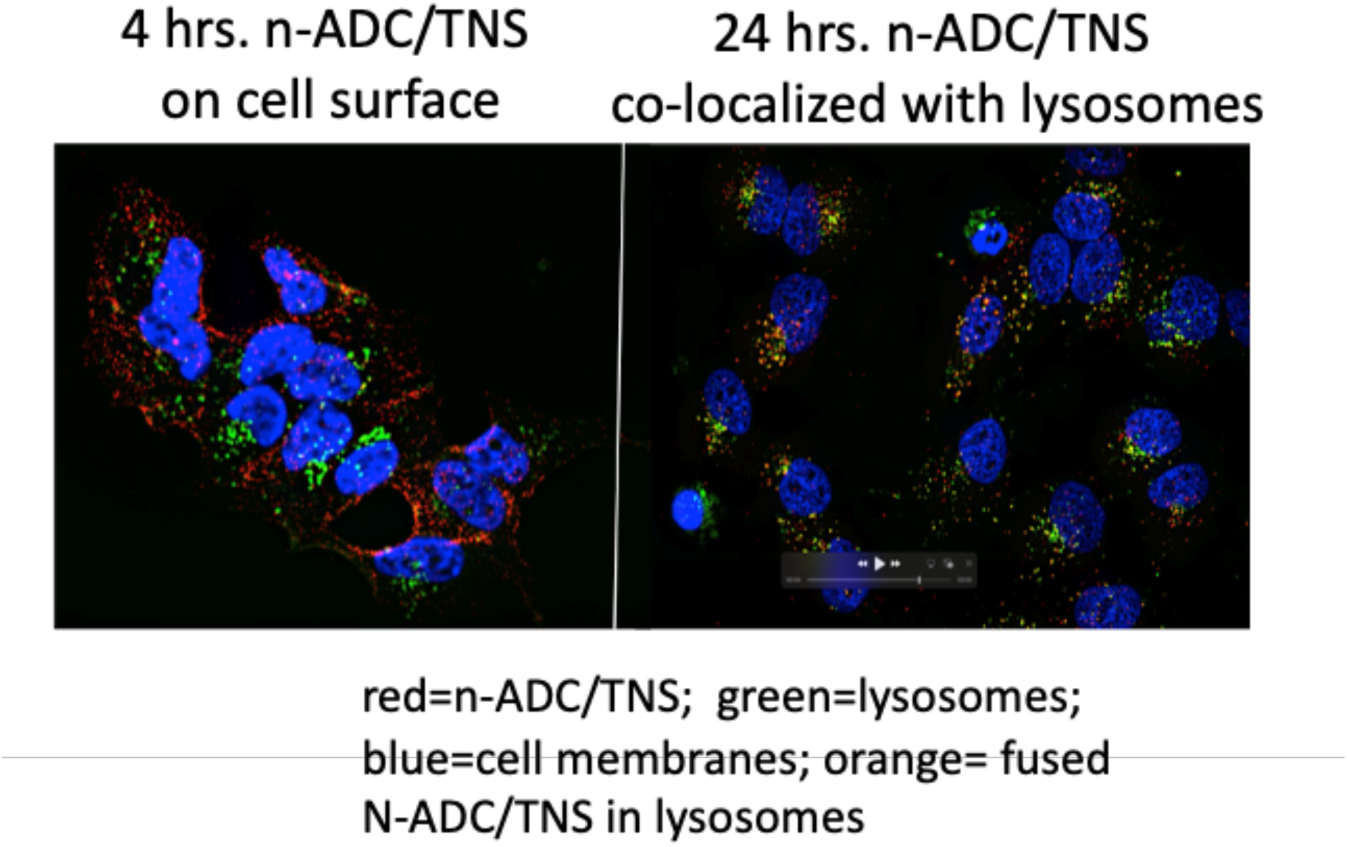
Red nanoparticles are seen bound to a Ewing tumor cell surface on the left after 4 hours incubation. At 24 hours they have entered the cell, fused with lysosomes (green) and the payload released, damaging nuclear DNA (blue) leading to chromatin disruption and cell death (right).

CD19, a type I transmembrane protein with a single transmembrane domain, is a B-cell-specific cell surface antigen and a key target for immunotherapy of B-cell malignancies and autoimmune diseases, overexpressed on the surface of most B-cell leukemias and lymphomas, including B-cell acute lymphoblastic leukemia (B-ALL) and chronic lymphocytic leukemia (CLL) [56]. At the same time, CD99 (also called MIC2), a transmembrane glycoprotein involved in cell adhesion, migration, and apoptosis, is overexpressed in various cancers, including ES [57]. In ES, the tumor cell surface antigen CD99 is a direct target of the ubiquitous EWS-ets fusion gene found in all true ESs and is thus universally expressed in ES. Human monoclonal antibodies with sufficient specificity to CD19 and CD99 are available. From a therapeutic delivery perspective, the candidate tumor biomarker should get internalized when bound to proteins at the cell surface [59, 59]. This allows targeted nanoparticles and, therefore, their therapeutic payload to enter tumor cells through receptor-mediated endocytosis.

### Tumor cell binding optimization

To determine the optimal amount of targeting antibody that gives the maximal cell binding, an increasing amount of antibody-conjugated micelle particles were added to fixed amounts of HPLNs. **Figure 5** shows the binding of *anti*-CD99-HPLN and *anti*-CD19-HPLN-binding to TC71 (ES) and REH (leukemia) cells by FACS, respectively, as a function of added micelle targeted with antibodies.

**Figure 5.**
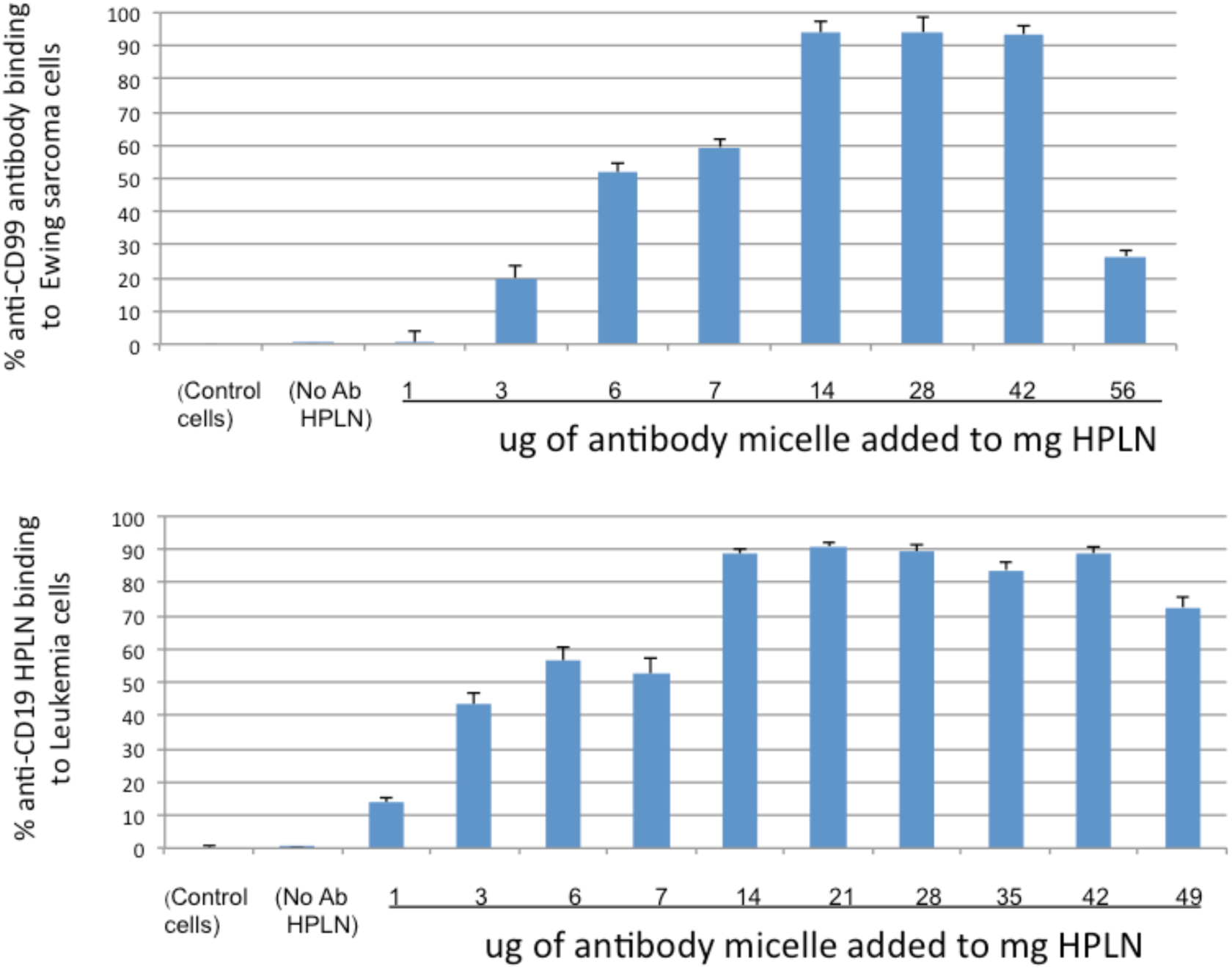
Human tumor cells were exposed to various test antibody-labeled HPLNs created by varying the ratio of Ab-micelles mixed with a fixed amount of HPLN. **A.** Human ES TC71 cell binding to *anti-*CD99 labeled HPLN, and **B.** Human leukemia REH cell binding to *anti-*CD19 labeled HPLN were assessed by FACS analysis using the innate HPLN fluorescence.

At low levels of antibody micelle to HPLN, the cell binding was minimal. As expected, increasing the amounts of Ab-micelle relative to HPLNs for both types of antibodies increased the cell affinity, and the FACS-measured binding described a bell-shaped curve. Interestingly, the binding to cells dropped off at the *high end* of the micelle to HPLN ratio, for either *anti*-CD19 or *anti*-CD99 antibody. One can speculate that beyond a specific surface density of antibody, an adverse binding effect comes into play, perhaps due to steric repulsion between HPLN and the cell.

For both *anti*-CD99 and *anti*-CD19 targeted HPLNs, we observed a maximum cell binding by FACS for about 14 to 42 μg antibody micelle mixed per mg of nanoparticle. Above this maximum, adding more antibody-conjugated micelle to HPLN solution showed a gradual attenuation of cell binding.

The final nADC/TNSs, NV101, NV102, and NV103, were tested in vitro for binding to either Ewing TC71 cells (NV101 and NV103) or Leukemia REH cells (NV102), by FACS. The results of binding time-course are reported in Supplementary Data.

### Induction of Apoptosis in Ewing Tumor Cells by NV101 versus Free Drug

The doxorubicin-loaded HPLNs were mixed with *anti*-CD99 conjugated antibody micelles to produce NV101. Binding of the empty *anti*-CD99 antibody labeled HPLNs to human ES (TC32) cells was confirmed by FACS analysis (see **Figure 5A**). Using the optimal Ab-micelle to HPLN ratio of 28 μg micelle to 1 mg HPLN/Dox, experiments were then performed to determine whether the NV101 antibody targeting function induced apoptosis. As shown in **Figure 6**, higher caspase-3 activity was observed in TC32 cells treated with NV101, which, in contrast, was not observed in the free drug doxorubicin or untargeted HPLN/Dox-treated cells. Moreover, for the well-known caspase-3 substrate poly (ADP-ribose) polymerase (PARP) [60], we observed marked cleavage of this substrate in the NV101-treated cells, whereas the free drug doxorubicin or untargeted HPLN/Dox-treated cells showed much lower levels of cleavage.

**Figure 6.**
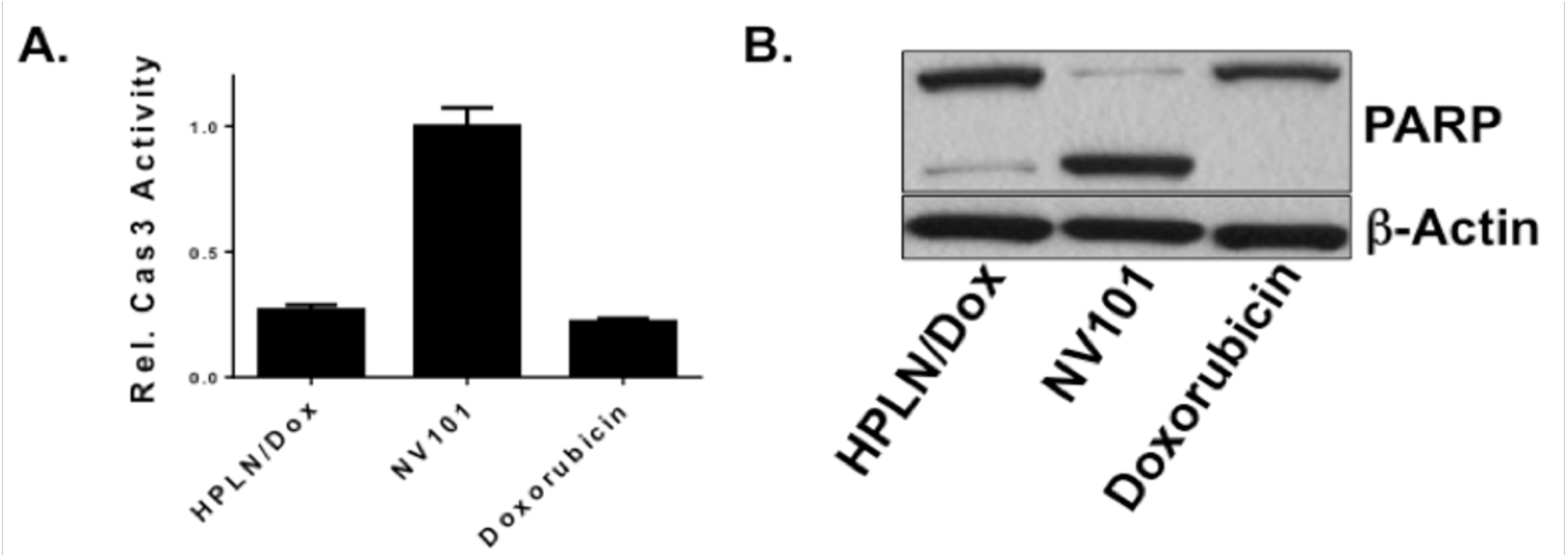
Induction of Apoptosis by NV101. TC32 cells were treated with one μM of doxorubicin, HPLN/Dox, or NV101 for two hours. Then the cells were washed with fresh media, followed by 24 hours of culture. A. Caspase-3 activity was measured in cell lysates by fluorimetry using caspase-3 cleavage kits, adjusting for protein concentration, where the non-treated cells were normalized to an arbitrary value of 1.0 B. Lysates from treated cells were assayed for PARP cleavage by Western blotting using antibodies to PARP. β-Actin was used as a loading control.

### Ewing Sarcoma Treatment with NV101

The efficacy of NV101 was evaluated using TC71 (ES) in a *NOD/SCID* mouse xenograft model. The TC71 cells were sourced from a patient with relapsed ES [61], whose tumor had developed drug resistance, rendering further treatment ineffective. This treatment-resistant, patient-derived tumor provided an ideal model to assess the potential of the targeted nanoparticle therapy. The TC71 cells were transfected to express the luciferase gene to enable monitoring of tumor burden by bio-luminescent (Xenogen) camera imaging [62]. HPLNs that were doxorubicin loaded were mixed with *anti*-CD99 conjugated antibody micelles to produce the nADC/TNS NV101. FACS analysis confirmed the binding of *anti*-CD99 antibody-labeled HPLNs to TC71 cells (**Figure 5A**).

Six *NOD/SCID* mice were injected with (1×10^6^) luciferase-transfected TC71 cells in the subcutaneous model. Six *NOD/SCID* mice were injected with (5×10^6^) luciferase-transfected TC71 cells in the metastatic model. Three received only buffer and three received NV101. The dosing schedule was conducted so that the animals received twice weekly at a dose of 2 mg/kg Dox. The tumor burden was assessed comparing the control animals with untargeted HPLN/Dox and NV101 over a 50-day time course.

The results are shown in **Figure 7**. After 20 days of treatment, there was a measurable difference in tumor burden between the NV101 treated and control group animals. This was seen in both the subcutaneous and metastatic Ewing tumor groups.

**Figure 7.**
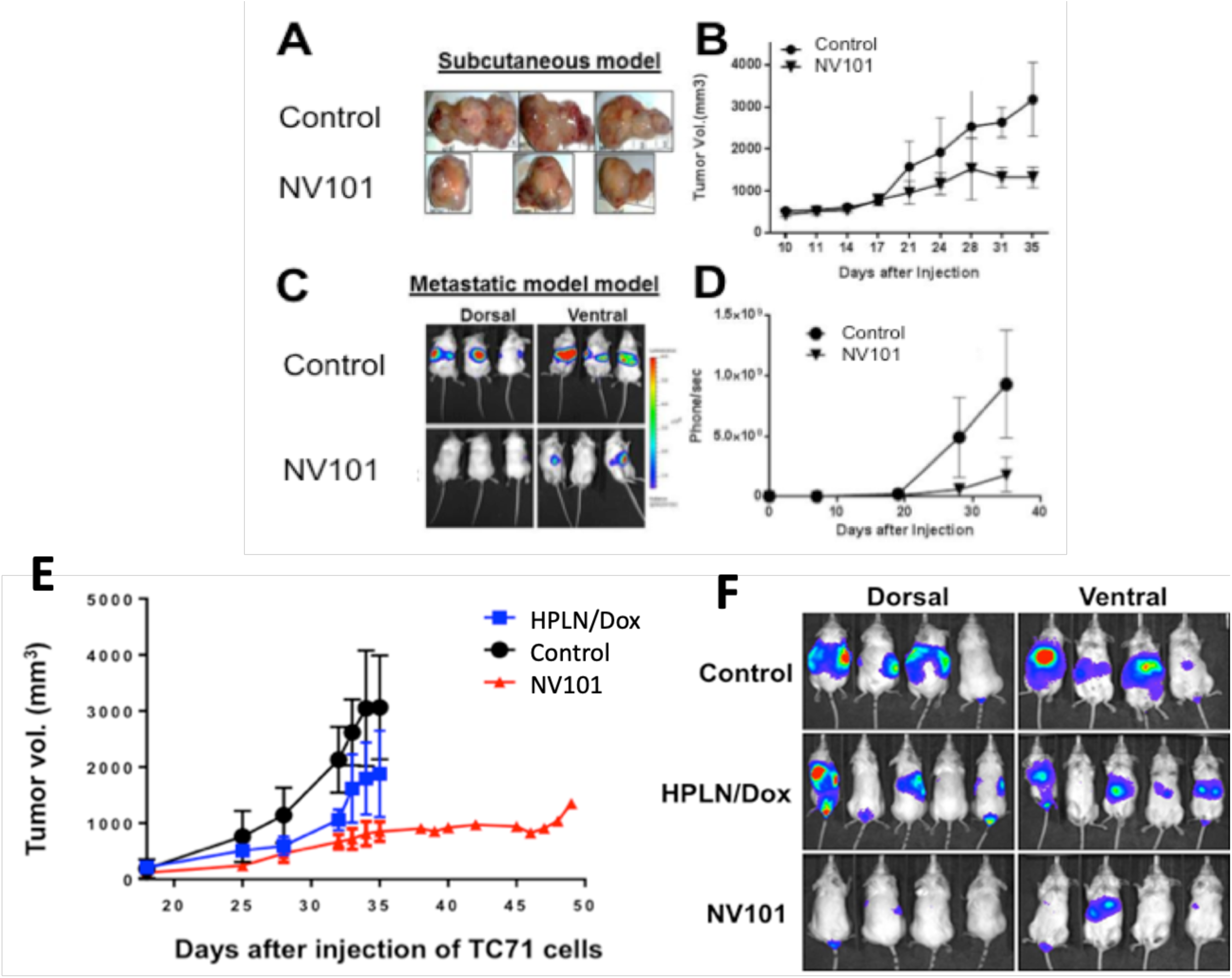
Subcutaneous model: (A) picture of an excised Ewing tumor removed from a control (untreated) animal (top row) compared to an NV101-treated animal (bottom row). (B). The tumor burden over the 35-day time course with twice-weekly NV101 treatments. Metastatic model: (C). Xenogen image from day-35 comparing the control tumor size (top row) with the NV101-treated animals. **D**. The tumor burden over the 35-day time course with twice-weekly NV101 treatments. **E.** Estimation of the TC71 tumor burden from the Xenogen camera images, over the treatment course from start of treatment on day 8, to day 50. **F.** Xenogen camera images of xenograft mice injected with patient-derived Ewing sarcoma cells (TC71) at day 24.

In Panel A, tumor burden estimation in caliper reveals a significant reduction in the NV101-treated group, whereas the control group exhibits continuous tumor growth. The untargeted HPLN/Dox group shows a moderate reduction in tumor burden, but it is less effective than NV101. In Panel B, Xenogen images of xenograft mice injected with TC71 cells on day 38 further confirm these findings, as the NV101-treated group displays a noticeably lower bioluminescent signal, indicating a smaller tumor burden. These results highlight the superior anti-tumor efficacy of NV101, making it a promising candidate for treating drug-resistant ES.

### Tumor Ablation in a transgenic mouse model of refractory human leukemia with NV102

Adult patient-derived ALL leukemia cells (*LAX7R* cells) were injected in *NOD/SCID* mice. *LAX7R* PDX cells are derived from an adult patient with relapsed treatment-resistant (i.e., refractory) ALL. *LAX7R* is BCR-ABL negative (“breakpoint cluster region-Ableson1”). The BCR-ABL fusion gene is found in 25-30% of adult ALL patients, and treatment can be successful with a variety of chemotherapeutic agents. However, BCR-ABL-*negative* cases are less likely to be responsive to these drugs [63]. Xenografted patient leukemic cells thus provided an excellent test case for our targeted nanoparticle therapy to demonstrate efficacy with a challenging, treatment-resistant, patient-derived tumor.

The *LAX7R* cells were transfected to express the luciferase gene to enable monitoring of tumor burden by bioluminescent camera imaging (*Xenogen^TM^ In Vivo Imaging System, IVIS*) [62]. We confirmed that this leukemia cell type expressed CD19 surface antigens and that NV102 bound to the *LAX7R* cells *in vitro* (data not shown). Fifteen *NOD/SCID* mice were injected with (5 x10^6^) luciferase-transfected *LAX7R* cells. Five NOD/SCID mice received only buffer, while five received doxorubicin-filled untargeted HPLNs, and five received NV102. The dosing schedule was conducted so that the animals received the same dose of doxorubicin (as HPLNs) twice weekly at a dose of 2 mg/kg doxorubicin. The results are shown in **Figure 8**.

**Figure 8.**
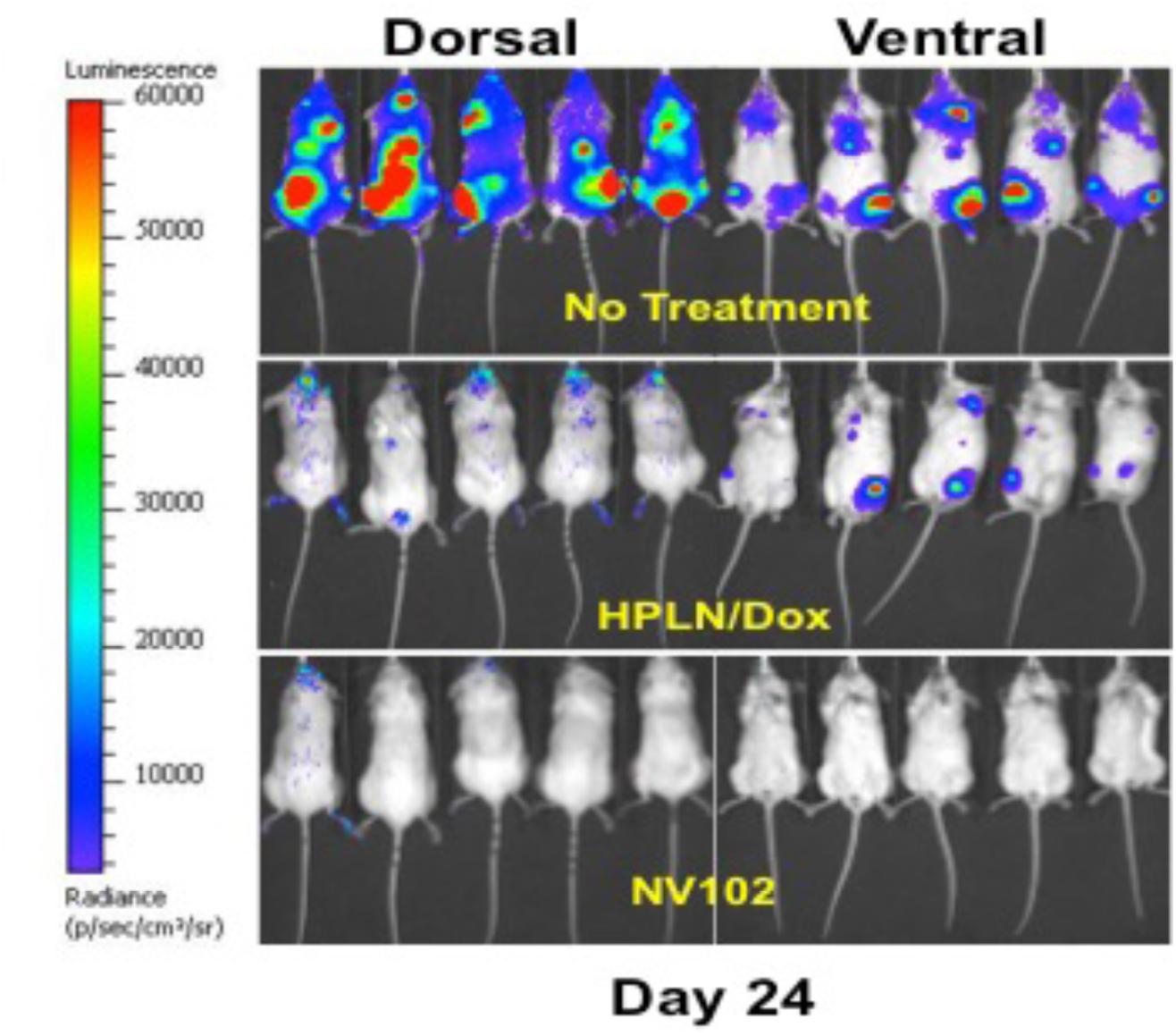
Xenogen camera images of xenograft mice injected with patient-derived ALL cells (LAX7R) at day 24.

After 24 days (16 days after the start of the first treatment) the observed tumor burden was extensive in the *buffer-only* control animals (top image). By day-38 the control animal tumor burden required euthanization. In contrast, the tumor burdens in the animals receiving either the *untargeted* HPLN/Dox (middle image) or NV102 (lower image) showed a dramatically reduced tumor burden.

The animals treated with *untargeted* HPLN/Dox showed a low tumor burden until day 55, after which they grew exponentially over the remaining three weeks of the experiment (**Figure 9A**). In these animals, the low tumor burden varied slightly from week to week up until day 55, and at this time point, the tumor growth broke out and they grew exponentially over the remaining three weeks of the experiment (**Figure 9B**). After another 10 days, the size of the tumors in these untargeted HPLN/Dox-treated animals quickly became as extensive as the buffer-only treated animals and required euthanization.

**Figure 9.**
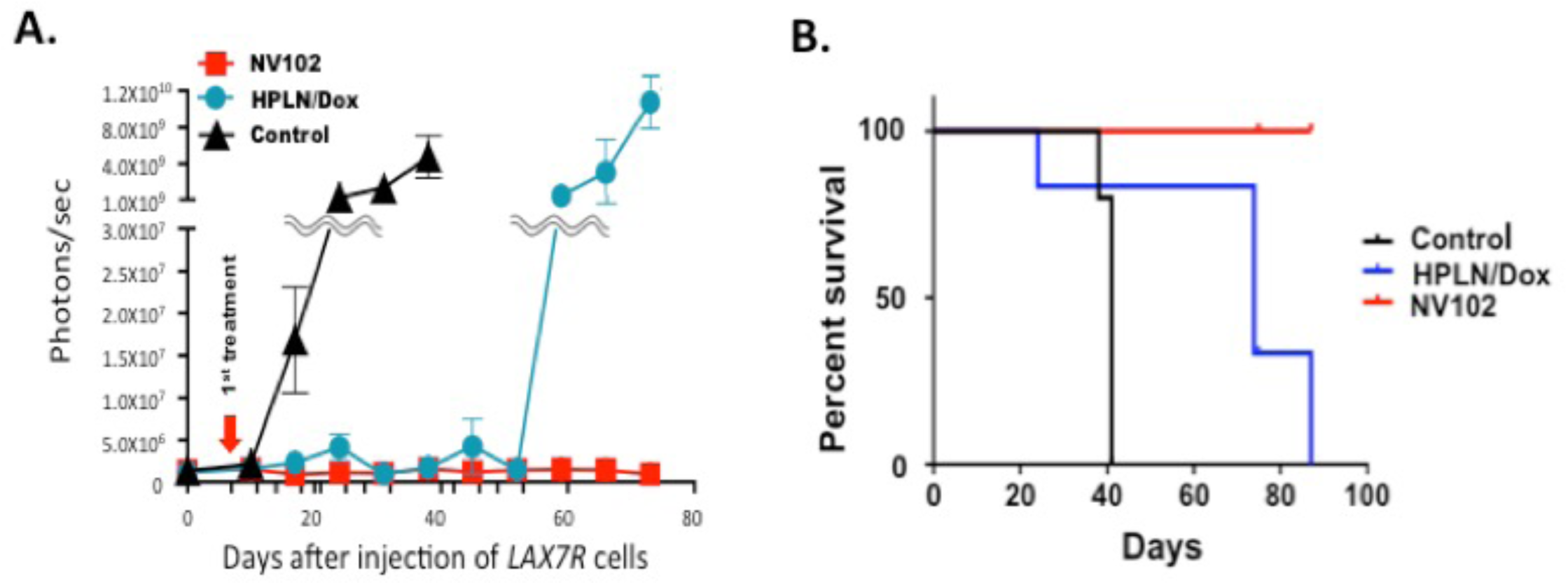
**A.** Estimation of the LAX7R tumor burden from the Xenogen camera images, over the treatment course from the start of treatment on day 8, to day 78. **B.** 87-day survival curve for the three groups of mice receiving human LAXR7 (ALL) leukemia cells treated with 1) buffer control, 2) untargeted HPLN/Dox, and 3) NV102.

Kaplan-Meier survival analysis is shown in **Figure 9B**. All NV102-treated animals survived and appeared healthy at day 87, when all control and *untargeted* HPLN/Dox-treated animals were dead of disease. After euthanization, necropsy revealed no evidence of disease in the NV102-treated animals.

### Treatment of Ewing Sarcoma cells with NV103 (irinotecan-containing nADC/TNS)

In a process similar to NV101, HPLNs were loaded with irinotecan (instead of doxorubicin) and were mixed with *anti*-CD99 conjugated antibody micelles to produce the nADC/TNS NV103. This study compares the tumor burden reduction in ES tumor-bearing mice with NV101, NV103 and untargeted NV101, to untreated control animals (5 per group). Eight-week-old female *NOG* mice (Taconic Hudson, NY, USA) were subcutaneously implanted on day 0 with 2 x 106 TC71-Luc Ewing cells. Seven days after the tumor cells were injected the mice were intravenously administered nanoparticles at a dose of 5 mg Ir/kg or 2 mg Dox/kg, via tail vein injection twice per week. As seen in **Figure 10** the control animals developed significant tumor burden, the Dox-containing NV101 had marginal tumor growth inhibition but the targeted, irinotecan-containing nanoparticles (i.e., NV103) resulted in complete tumor ablation.

**Figure 10.**
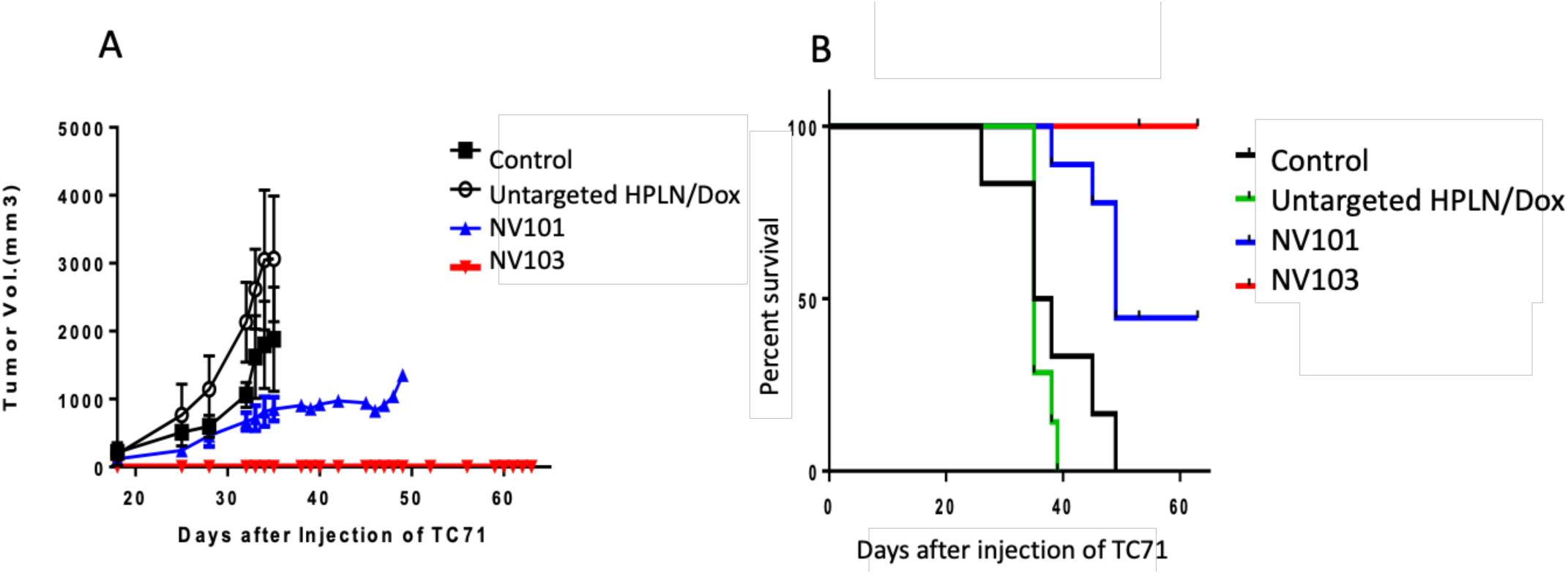
**A**. Estimation of the ES tumor burden from the Xenogen camera images, over the treatment course from the start of treatment on day 8, to day 65. **B.** 65-day survival curve for the three groups of mice receiving human TC71 cells treated with 1) buffer control, 2) untargeted HPLN/Dox, 3) NV101 and 4) NV103.

### Determination of actual number of antibodies per NV103 particle and irinotecan molecules per antibody

Using a NanoSight^TM^ (NS300, Malvern Panalytical) instrument for quantification, we observed that a 1 mg/mL NV103 concentration is equivalent to 7.22E+13 particles per mL. We analyzed the actual amount of antibody protein per mg of HPLN by ELISA. This translates to about 8 antibodies per individual HPLN particle. Analysis of the irinotecan content in one NV103 particle revealed 10,000-20,000 Irinotecan molecules per particle. This translates to 1,250 to 2,500 irinotecan molecules delivered to the target cell *per antibody,* compared to 2-8 drug molecules per antibody reported for the typical ADC [6].

## DISCUSSION

Liposomes, unilamellar vesicles composed of natural and/or synthetic lipids, have been intensively studied as drug delivery vehicles [64]. One significant advantage of a liposomal delivery vehicle is the *high payload carrying capacity*. This allows them to carry high concentrations of small molecules without exposure to plasma. The membrane shell provides a surface onto which a targeting moiety can be attached. The therapeutic potential of cancer chemotherapy is often restricted by dose-limiting toxicities affecting bone marrow, GI tract, liver, heart, and other tissues. These side effects determine the *maximum tolerated dose* (MTD) of the drug that can be safely delivered without precipitating unacceptable adverse events. Therefore, complete ablation of primary or metastatic tumor with chemotherapeutics alone is rarely achieved, although if higher doses were tolerable, complete ablation would be possible [65].

Significant improvements in survival and/or efficacy have not yet been realized in the treatment of many malignancies for several decades, despite the use of new drug combinations. On the contrary, intensified therapy has instead resulted in treatment-induced morbidity and secondary malignancies in pediatric patients after otherwise curative therapy [66]. Targeted forms of therapy like ADCs have been developed and are increasingly used in an attempt to circumvent this issue by directly targeting therapeutic drug molecules to tumor cells [67]. However, since the drug is covalently linked to the antibody, it is not bioavailable until released, typically through scission of a linker. This stepwise process requires the ADC to remain intact in circulation until it reaches the intended target, and subsequently, after internalization, release the drug inside the target cells. ADCs are also limited by the number of drug molecules that can be attached per antibody (typically a drug-to-antibody ratio or DAR of 2 to 8), and attempts to increase the drug-protein ratio often cause antibody deactivation [68]. Long-term treatment using ADCs can also induce an immune response to the targeting antibody. Thus, while ADCs address some shortcomings of traditional chemotherapy, several technical challenges have limited their efficacy [69].

Prior efforts using drug molecules encapsulated in *targeted* nanoscale carriers have not yet resulted in complete tumor ablation. This failure is more a result of tumor cell antigen target, targeting agents, and/or tumor types [70,71]. The tumor microenvironment also hampers targeted nanoparticles in cancer treatment because tumor stroma impedes nanoparticle access to tumor cells. Failures to choose a suitable tumor system, a tumor antigen unique to that tumor, a suitable targeting antibody and an optimal drug has led to recent lack of success [72].

As a technology, utilizing tumor-specific targeting agents on the surface of drug-encapsulated nanoparticles can: (a) ***preferentially deliver*** the drug to cancer cells while sparing normal cells and tissues, and (b) deliver lower ***systemic*** cytotoxic doses to minimize both initial and long-term toxicity and morbidity or mortality. Ideally, the drug should be delivered preferentially to the tumor at high dose levels and with minimal normal, or healthy, tissue exposure.

The nADC/TNP platform is a unique and versatile drug delivery technology helpful to treat a variety of cancers and potentially other diseased tissues as well. The hollow shell particle is amenable to the addition of various targeting agents such as antibodies [4], diabodies [46], and peptides [73]. Once polymerized, the nanoparticle is loadable with cytotoxic drugs. The advantage of a liposomal delivery vehicle is the high payload carrying capacity. Although our nADC/TNPs were initially formulated for their stable drug loading characteristics, they also proved to be more therapeutically potent than their conventional liposome analogs. Once inside the target tumor cell, the cytotoxic drug in the nADC/TNP is released in the lysosome and diffuses to the nucleus, causing irreparable DNA damage and tumor cell death. This boost in potency appears to directly depend on the polymerizable PCDA lipid content [46].

nADC/TNSs provide a greater on-target drug to antibody ratio than an ADC; a *thousand* drug molecules per nanoparticle-conjugated antibody versus only 2 to 8 drug molecules per ADC antibody [68]. In addition, the polyvalency of many antibodies attached to a single nanoparticle enhances the overall avidity of nanoparticles to target tumor cells, compared to a single ADC molecule. Furthermore, encapsulation limits plasma concentrations of free drug and exposure to normal tissues.

And lastly, in a hollow-shell nanoparticle, the drugs are not chemically conjugated to the carrier, as they are in the ADC assemblies. Once released from the carrier, no *in situ* chemical decoupling is required for the drug to become fully bioavailable.

nADC/TNS builds on the advantages of conventional liposomal nanoparticles by creating a more stable nanoparticle that is less prone to leakage, which also imparts a favorable route of uptake via endocytosis properties that are not true of all liposomal nanoparticles. The use of fully polymerized or partially polymerized vesicles has been described and studied for use in diagnostic and cancer therapies for almost 30 years [7]. Unlike conventional liposomes, HPLNs can also be made to be intrinsically fluorescent. Upon initial polymerization, the “blue” form of the polydiacetylene polymer is non-fluorescent. Heat treatment of the blue vesicles leads to a color change to red-pink and the formation of a fluorescent fluorophore [44]. The color change is a result of a subtle conformational change in the polymer backbone, leading to the fluorescent polymer form. The optical fluorescence property is a side benefit that allows the HPLNs and subsequent nADC/TNSs to be readily traced from the time they bind to target cells until they are deposited and compartmentalized into subcellular structures. A unique property of the polydiacetylene fluorescence is that little or no photobleaching occurs. This, in turn, enables semi-quantitative image analysis measurements of nanoparticle concentrations in tumor and normal tissues.

Partial polymerization of nanoparticles for drug delivery is unique and innovative, and the physiological consequences in biological systems are still being investigated. For nanocontainers (nanoparticle-based) drug encapsulation, the problem of containment versus controlled release of anti-cancer agents has long been a challenge. On the one hand, nanoparticles need to be formulated to allow for efficient packaging of therapeutic agents and stable containment of drug post-manufacture through administration, while on the other hand, once they have localized to target tissues or tumors, they need to be able to facilely release their payload to have the desired therapeutic effect. This latter attribute has been particularly challenging to design into conventional liposome formulations [74]. The prior efficacy studies suggest that the presence of the polymer in the HPLN should enhance the stability of the encapsulated drug towards passive leakage [48–51] while also enhancing the bioavailability of the drug [46, 49, 50], when compared to the conventional liposome forms. It could also promote endocytic uptake and lysosomal payload release of nADC/TNSs, as opposed to cell membrane fusion that is typical of many liposomal nanoparticles, which results in direct exposure of the drug payload to drug efflux pumps like MDR1 [3].

The observation of an enhanced drug release accentuated under acidic conditions suggests that the superior HPLN/drug efficacy, compared to conventional liposomes, may be due to more facile HPLN particle opening and drug release in the lower pH physiological environment that would be expected from receptor-mediated endocytic conditions in late endosomes and lysosomes [75]. We therefore hypothesize that the more efficient tumor cell killing *in vitro* by the HPLN is a direct result of the labile nature of the polymer-containing membrane, specifically under acidic conditions. This suggests that *any* drug delivered to a tumor cell in an HPLN will be more efficacious than a drug delivered by a conventional liposome formulation.

Our goal in creating the three formulations discussed in this paper was to document the broad utility of nADC/TNSs across multiple payloads and antibodies against a selection of suitable tumor types. The results document the desirable carrying and drug-loading properties of HPLNs that also impart additional stability by partially polymerizing the membrane of nADC/TNSs. To this, we have further incorporated a surface-attached, covalently linked antibody that recognizes an over-expressed tumor cell protein that is absent from most (CD19) or nearly all (CD99) normal tissues, thereby enabling preferential drug delivery to the tumor while protecting normal tissue from similar doses of the drug. The nADC/TNS platform is versatile enough to encapsulate different kinds of cancer chemotherapeutic drugs and vary the targeting agent to customize particle delivery to different tumor types, thus enabling optimal treatment of a wide variety of tumor types. For our initial studies, we chose the drugs irinotecan and doxorubicin and antibodies targeting CD19 (acute lymphoblastic leukemia and other hematopoietic malignancies) and CD99 (ES).

Acute lymphoblastic leukemia (ALL) is a hematologic malignant disorder originating from either T- or B-cell lymphoid precursor. ALL is the most common childhood leukemia and, in adults, the third most common leukemia. ALL patients are generally treated with chemotherapeutic agents including doxorubicin, vincristine, prednisone, cyclophosphamide, and asparaginase [76]. While effective for most pediatric ALL patients, this multi-agent chemotherapy protocol has been far less successful with adult patients. These *LAX7R*-cells used here were derived from a patient whose leukemic cells were unsuccessfully treated with chemotherapy [77]. Other xenograft models of B-ALL have been treated with dexamethasone, vincristine and doxorubicin [78]. Treatment of these cells by NV102 showed dramatic tumor burden reduction in the mouse model. This result implies that even single-agent treatment with NV102 tumor-targeted therapy is more effective than even combinatorial chemotherapy that includes the same drug. It also suggests that nADC/TNSs containing other agents like the ones used in this protocol would further enhance the efficacy of this therapeutic approach.

While *in vitro,* the difference in tumor cell killing between *untargeted* HPLN/Dox and NV102 was small (see Supplementary data section), the adult treatment-resistant ALL leukemia cells in *NOG/SCID* mice showed a dramatic response over 87 days of treatment (**Figure 9**). In contrast, the tumor burden in mice treated with *untargeted* HPLN/Dox remained low, but after day 55 the tumor size increased and became equal to that of control animals by day 87. On the other hand, the NV102-treated animals had no detectable tumor burden through day 87, resulting in 100% survival. This is a remarkable result, given the known treatment resistance of these leukemic cells derived from an adult patient who did not respond to similar drugs as free drugs (not encapsulated in nanoparticles).

ES-patients will generally be treated with combination chemotherapy, consisting of three different topoisomerase inhibitors out of seven drugs used for front-line and relapse therapy (e.g., doxorubicin, etoposide, and irinotecan) [79]. This is necessary because of the overall poor prognosis and historical inability to cure the tumor with surgery or radiation therapy alone. Only systemic multi-agent therapy has resulted in cures, and the cure rate (∼50%) has not changed appreciably in decades. In a search for drugs showing particular efficacy in the treatment of ES to improve the static survival rate, Irinotecan was found to be particularly effective [57]. This effect is linked to high expression of a DNA/RNA helicase-related gene, SLFN1, that is expressed at high levels in ES, which increases sensitivity to this drug [57]. Irinotecan is already used in combination with temozolomide for the treatment of relapsed ES patients, though it is not clear that such treatment enhances survival. Irinotecan is unique among approved commercial cytotoxic agents specifically because it is a pro-drug requiring carboxylesterase-2 (CE-2) based enzymatic conversion to the active metabolite SN-38. SN-38 is 100 to 1000 times more cytotoxic than the irinotecan pro-drug form, which prevents its use as a free drug in humans [80]. However, ES cells express high levels of the CE-2 enzyme, resulting in rapid conversion of irinotecan to SN-38 in tumor cells, which is extremely toxic to Ewing tumor cells. Comparison of the two CD99 targeted nADC/TNSs NV101 and NV103 in mice with implanted human ES cells, showed the irinotecan encapsulated formulation was more efficacious than the doxorubicin formulation (**Figure 10**). This observation of superior tumor kill in mice will be characterized in greater detail in a future publication.

In measuring the toxicological and PK/PD parameters for NV103 in normal mice (see Supplementary data section) the most striking observation from this investigation was that while the free Ir injection is given at a **10-fold greater dose than the Ir contained in the NV103 particles**, the plasma half-life, Cmax and AUC measurements all show a dramatically higher amount of Ir in the plasma of the NV103 treated mice. While the time to establish the maximal Ir concentration (Tmax) is similar for both NV103 and free Ir, the clearance of Ir described as the amount remaining in circulation (AUC) suggests that the circulation kinetics are greatly affected by the form administered (encapsulated vs. free), with the encapsulated form showing more than 5-fold increased AUC verses free Ir. Another analysis looked at the plasma levels of the active metabolite SN-38. SN-38 plasma levels for NV103 treated animals resulted in a greater Cmax and longer circulation half-life than in animals with free Ir. The final PK analysis assessed the Ir bone marrow concentration versus time profile. In contrast to the plasma Ir levels, the Ir derived from NV103 compartmentalized to bone marrow **4-fold less** than free Ir. Bone marrow toxicity is a major limitation for cancer chemotherapies, and NV103 encapsulation appears to significantly reduce this limitation.

## CONCLUSIONS

*In vitro*, the addition of antibody targeting provided a greater cytotoxicity boost for NV101 over untargeted HPLN/Dox (60%) at the 24-hour time point. NV102 showed a small gain in tumor cytotoxicity over untargeted HPLN/Dox (11%) at the 24-hour time point. NV103 was 25-53% more effective compared to HPLN/Ir, at all time points. Overall, targeting uniformly enhances tumor cell kill compared to untargeted counterparts (See Supplementary data section).

*In vivo*, NV101 demonstrated tumor burden reduction in *NOG/SCID* mice with growing ES tumors in both subcutaneous flank tumors and in a metastatic tumor model compared to control cells and untargeted HPLN/Dox. NV101 activity is due to Dox-induced DNA double-strand breaks causing irreparable DNA damage, leading to apoptosis, as documented by high caspase-3 activity. The time course study showed tumor size that plateaued by day-35 and continued to day-50, whereas all the control and non-targeted HPLN/Dox treated animals were sacrificed due to rapidly growing tumors whose size exceeded those allowed by the animal control committee.

In ALL, NV102 showed dramatic tumor burden reduction resulting in 100% survival in the treated mice. The treatment of a human PDX model of relapsed, treatment-resistant adult ALL dramatically shows the benefit of antibody targeting over untargeted particles.

Comparing NV101 and NV103 in *NOG/SCID* mice with growing ES tumors, greater efficacy has been demonstrated, using irinotecan-containing nanoparticles (NV103) over doxorubicin nanoparticles (NV101). Only NV103 demonstrated complete tumor eradication and 100% survival (**Figure 10**).

While pre-clinical use data, as reported here, is limited to a proof-of-principle studies using drug-containing targeted nADC/TNSs, the preliminary data are very encouraging. Studies *in vivo* with ES, high-grade glioma (GBM), and a variety of other typical adult solid tumors, including triple-negative breast cancer (TNBC), prostate, colon, lung, pancreatic, and liver cancer, are also underway. The outcomes of these studies will be reported in detail in subsequent publications.

## Materials and Methods

### Reagents

HPLN and micelles are comprised of hydrogenated soy L-α-phosphatidylcholine (hydro-soy-PC, Nanosoft Polymers, Winston-Salem, NC), 1,2-disteroyl-*sn*-glycero-3-phosphoethanolamine-N-[methoxy(polyethylene glycol)-2000] ammonium salt (m-Peg2000-DSPE), cholesterol (ovine wool), 1,2-disteroyl-sn-glycero-3-phosphoethanolamine-N-[maleimide(polyethylene glycol)-2000] ammonium salt (mal-Peg2000-DSPE) all from Avanti Polar Lipids (Alabaster, AL). PCDA lipids: N-(methoxy(polyethylene glycol)-2000)-10-12-pentacosadiynamide (m-Peg_2000_-PCDA), and N-(5’-hydroxy-3’-oxypentyl)-10-12-pentacosadiynamide (h-Peg_1_-PCDA) are prepared at NanoValent, synthesis described below. The sources of the monoclonal antibodies were *anti*-CD99 human antibody (Curia, Belmont, CA) and *anti*-CD19 human antibody (Creative Biolabs, Shirley, NY). Other drugs and reagents: doxorubicin hydrochloride salt (LC Labs, Woburn, MA), and irinotecan hydrochloride salt (Thymoorgan Pharmazie Gmbh, Germany), Triton X-100 (Amresco, VWR, Radnor, PA). Ferric chloride, ammonium thiocyanate, phosphate buffered saline (PBS, pH 7.4), ammonium sulfate, chloroform, magnesium sulfate (anhydrous) ethyl acetate, hexane, methanol, methylene chloride, L-cysteine hydrochloride monohydrate (cys), tris(2-carboxyethyl) phosphine hydrochloride (TCEP), thiazolyl blue tetrazolium bromide, N-hydroxysuccinimide, 10,12-pentacosadiynoic acid (PCDA), 1-(3-dimethylaminopropyl)-3-ethylcarbodiimide hydrochloride (EDC), 2-(2-aminoethoxy) ethanol, triethyl amine, all from MilliporeSigma, (Burlington, MA). Mpeg45-NH2 is purchased from PurePEG Cat. #364645 (San Diego, CA).

### N-succinimidyl-10, 12-pentacosadiynate

10,12-pentacosadiynoic acid (PCDA) is dissolved in chloroform and filtered to remove any blue polymer followed by recrystallization from hot 95% ethanol. After evaporation and drying under vacuum, solid PCDA (1.00 g, 2.7 mmol) was dissolved in CH_2_Cl_2_ (10 mL) and N-hydroxysuccinimide (.345 g, 3.0 mmol) was added followed by 1-(3-dimethylaminopropyl)-3-ethylcarbodiimide hydrochloride (EDC) (.596 g, 3.1 mmol). The progress of the reaction was followed by silica gel TLC with chloroform:methanol (20:1) as eluent. The solution was stirred at room temperature for two hours followed by rotary evaporation of the CH_2_CI_2_. The residue was dissolved in a mixture of ethyl acetate:hexane (50:50) and water. After separation the aqueous layer was extracted three times with ethyl acetate:hexane (50:50) and the combined organic extracts washed with 0.5N HCl (3x), saturated aqueous sodium bicarbonate (1x) and finally saturated aqueous sodium chloride (1x) followed by drying with MgSO_4_. After filtration, the solvent was removed by rotary evaporation to give 1.21 g (2.6 mmol) of a crude white solid. The solid was recrystallized from hot absolute ethanol yielding 1.1g (85% yield). ^1^H NMR (500 MHz, CDCl_3_) δ 0.88 (t, 3H, J=6.7 Hz), 1.23 (br s, 26H), 1.50 (m, 4H), 1.71 (m, 2H), 2.21 (t, 3H, J=6.8 Hz), 2.56 (t, 3H, J=7.7 Hz), 2.81 (d, 2H, J=2.9 Hz); 13C NMR (CDCI3) δ 14.02, 19.06, 19.08, 22.58, 24.44, 25.49, 28.17, 28.25, 28.58, 28.68, 28.75, 28.79, 29.00, 29,24, 29.37, 29.50, 29.52, 29.54, 30.80, 31.81, 65.14, 65.23, 77.32, 77.46, 168.54, 169.11; HRMS FAB m/z for C_29_H_45_NO_4_ calcd 471.3349 (M+), found 471.3368.

### N-(methoxy(polyethylene glycol)-2000)-10-12-pentacosadiynamide (m-Peg_2000_-PCDA) <u>1</u>

A solution of 1.21 g (2.6 mmol) of N-succinimidyl-10,12-pentacosadiynate in 50 mL of CHCl_3_ was cooled in an ice bath. To this solution was added 0.30g (2.9 mmol) of 2-(2-aminoethoxy) ethanol followed and 0.36 mL (2.6mmol) of triethyl amine, and the reaction solution was allowed to warm to ambient temperature and stir overnight. The progress of the reaction was followed by silica gel TLC with chloroform:methanol (6:1) as eluent The reaction mixture was diluted with 25% isopropanol in chloroform, and washed with 0.5N HCl (3x), saturated aqueous sodium bicarbonate (1x) and finally saturated aqueous sodium chloride (1x) followed by drying with MgSO_4_. After filtration the solvent was removed by rotary evaporation, and the crude residue was purified by silica gel column chromatography eluting with a 10:1 chloroform:methanol solution. Isolate 0.77g of recrystallized product (64% yield). ^1^H NMR (500 MHz, CDCl_3_) δ 0.4–0.6 (t, 3H, J=11.6 Hz), 1.2-1.35 (m, 22H), 1.37-1.45 (m, 4H), 1.48-1.56 (m, 4H), 1.6-1.66 (bt, 2H, J=11.7 Hz) 2.14-2.21 (t, 2H, J=12.5 Hz), 2.24-2.28 (t, 4H, J=11.6 Hz), 3.4 (s, 3H), 3.44-3.48 (q, 2H, J= 8.6 Hz), 3.53-3.59 (m, 6H), 3.6-3.74 (bs, ∼170H), 3.76-3.79 (bt. 2H, J=8.1 Hz), 6.2 (s, 1H). HRMS MALDI m/z for C_116_H_225_NO_46_ calcd 2368.530 (M+), found [M+Na] = 2391.248.

### N-(5’-hydroxy-3’-oxypentyl)-10-12-pentacosadiynamide (h-Peg_1_-PCDA) *(2)*

A solution of 1.21 g (2.6 mmol) of N-succinimidyl-10,12-pentacosadiynate in 50 mL of CHCl_3_ was cooled in an ice bath. To this solution was added 0.30g (2.9 mmol) of 2-(2-aminoethoxy) ethanol followed and 0.36 mL (2.6mmol) of triethyl amine, and the reaction solution was allowed to warm to ambient temperature and stir overnight. The progress of the reaction was followed by silica gel TLC with chloroform:methanol (6:1) as eluent. The reaction mixture was diluted with chloroform, and washed with 0.5N HCl (3x), saturated aqueous sodium bicarbonate (1x) and finally saturated aqueous sodium chloride (1x) followed by drying with MgSO_4_. After filtration the solvent was removed by rotary evaporation, and the crude residue was recrystallized from hot chloroform:hexane (1:1.5) followed by silica gel column chromatography eluting with a 10:1 chloroform:methanol solution. Isolate 0.77g of recrystallized product (64% yield). ^1^H NMR (300 MHz CDCl_3_) δ 0.85 (t, 3H, J=6.5 Hz), 1.23 (br s, 26H), 1.50 (m, 4H), 1.71 (m, 2H), 2.14 (t, 2H, J=7.8 Hz), 2.18 (t, 4H, J=7.8 Hz), 2.4 (br s, 1H), 3.45 (t, 2H, J=5.0 Hz), 3.55 (m, 4H, J=6.0 Hz), 3.71 (br s, 2H), 6.0 (br s, 1H); 13C NMR (CDCI3) δ 14.02, 19.18, 19.21, 22.69, 25.68, 28.29, 28.36, 28.77, 28.86, 28.93, 29.10, 29.17, 29.23, 29.35, 29.48, 29.61, 29.63, 29.65, 31.92, 36.74, 39.20, 61.77, 65.21, 65.29, 70.00, 72.21, 77.46, 77.62, 173.37. HRMS ESI m/z for C_29_H_51_NO_3_ calcd 461.3861 (M+), found 462.3933 (M+H).

### HPLN preparation

A solution of lipids hydro-soy-PC, h-Peg_1_-PCDA, m-Peg2000-DSPE and m-Peg_2000_-PCDA and cholesterol in a molar ratio: 51:14:2:1:32, respectively were evaporated from chloroform to a thin film under high vacuum, overnight at ambient temperature. A 155 mM ammonium sulfate solution (pH 5.5) was added to the waxy solid lipid mixture to give a 10 mg/mL lipid and cholesterol solution and this was bath sonicated (VWR Ultrasonic Cleaner Mod 250D) for 30 minutes between 70-80°C. The solution was allowed to rest at ambient temperature for 30 minutes and then the bath sonication (70-80°C) was repeated for another 30 minutes. The resulting solution was nearly transparent with no visible or suspended solids. This solution was filtered through a sterile 0.45 µm Supor Membrane syringe filter, 32 mm Acrodisc (Pall Life Sciences, Port Washington, NY) and extruded at 50-55°C through a single, 47 mm, 80 nm pore size polycarbonate membrane (Whatman Nuclepore Track-Etch membrane, Cytiva Marlborough, MA) on a high-pressure extruder (Avestin, Emulsiflex-C5 homogenizer),10 times followed by 10 minutes of continuous extrusion. The DLS particle size was determined in ammonium sulfate solution on a Zetasizer Nano-ZS instrument (Malvern, Westborough, MA). The liposome solution was cooled overnight at 4°C. After warming to ambient temperature, the stirred solution was irradiated with UV light (254nm, Spectrolinker XL-1000 UV crosslinker, Spectronics Corp.) with constant stirring for 60 seconds to give a deep blue HPLN solution. The ammonium sulfate solution exterior to the HPLNs was removed and buffer shifted to PBS (pH 7.4) on a Millipore Labscale TFF system (#29751, MilliporeSigma, Burlington, MA) with a Pellicon XL Cassette (Biomax 1000 kDA, MilliporeSigma, Burlington, MA). The buffer was shifted by doing four 10X PBS exchanges. The HPLN concentration was determined by the Ammonium Ferrothiocyanate Assay (2.7 g ferric chloride, 3.0 g ammonium thiocyanate in 100 mL H_2_O). 80 uL of HPLN solution was added to 1.2 mL chloroform and 1.2 mL of ammonium ferrothiocyanate solution in a glass tube and shaken vigorously for 5 min on a Multi-Therm Shaker (Benchmark model H5000-H) at 1500 rpm. The tubes were spun on a bench-top centrifuge (Marathon 6K centrifuge) at ∼900 rpm for 2 min. The orange chloroform (lower) layer was transferred to a 1 cm path length 1 mL quartz Bio-cell chamber filled completely and read absorbance at 488 nm (Bio-Tek Synergy HT plate reader, PMT 49984). The HPLN concentration was calculated by: [HPLN] (mg/mL) = ((Abs_488_/0.3948)-0.0919) as established by a standard curve.

### HPLN/Dox

Solid doxorubicin hydrochloride (Dox) was added to a solution of HPLN in PBS at a concentration of 5mg/mL (2.5 mg Dox: 5 mg HPLN) and shaken at 60°C for 40 minutes. The solution was centrifuged (Thermo IEC Multispeed Centrifuge) at 10,000 rpm for 5 min. The orange HPLN/Dox supernatant was applied to the Millipore TFF system fitted with a Biomax 1000 kDA Pellicon XL cassette to remove the unencapsulated Dox. The unencapsulated drug was removed by doing four 10X PBS exchanges. The HPLN/Dox was centrifuged at 10,000 rpm for 15 minutes. The supernatant was filtered through a sterile 0.2 µm Supor Membrane syringe filter, 32 mm Acrodisc (Pall Life Sciences, Port Washington, NY). The particle size was assessed on a Zetasizer Nano-ZS. The HPLN/Dox particle concentration was determined by the Ammonium Ferrothiocyanate Assay previously described. The Dox concentration inside the HPLN is determined colorimetrically by rupturing the HPLN with 1% triton-X 100 and exposing the solution to sodium hydroxide. In microtiter well (A), 50 uL of HPLN/Dox was added to 100 uL PBS plus 110 uL DI water. In well (B), 50 uL of HPLN/Dox was added to 100 uL PBS plus 100 uL of 125 mM sodium hydroxide plus 10 uL 1% triton-X 100 (both solutions in DI water). Particle rupture was carried out by thoroughly mixing and allowing the reaction to occur at ambient temperature for 20 min protected from light. An absorbance was taken at 590 nm for both wells. The concentration of Dox is determined by taking the difference between well A and B and comparing to a standard curve. Loading efficiency can be determined by taking the Dox concentration in mg/ml divided by the HPLN concentration in mg/ml to yield mg Dox/ mg HPLN.

### HPLN/Ir

Irinotecan hydrochloride solution (20 mg/mL Ir HCL-trihydrate) was added to a solution of HPLN in PBS at a concentration of 5mg/mL for a final ratio of 5 mg HPLN to 5 mg of Ir, and shaken at 60°C for 40 minutes. The solution was centrifuged at 10,000 rpm for 5 min. The red HPLN/Ir supernatant was applied to the Millipore TFF system, fitted with a Biomax 1000 kDA Pellicon XL cassette, to remove the unencapsulated Ir. The unencapsulated drug was removed by doing four 10X PBS exchanges. The HPLN/Ir was centrifuged at 10,000 rpm for 15 minutes. The supernatant was filtered through a 0.2 u syringe filter. The particle size was assessed on a Zetasizer Nano-ZS. The HPLN/Ir particle concentration is determined by the Ammonium Ferrothiocyanate Assay previously described. The HPLN/Ir irinotecan concentration is determined by rupturing a 980 uL aliquot with 20 uL of 1% triton-X 100 (solution in DI water) for 30 min at ambient temperature. The solution is centrifuged at 10,000 rpm for 5 min. The absorbance of the supernatant is read at 370 nm. The Ir concentration is determined by comparing the 370 nm absorbance to a standard curve generated from the 20 mg/ml stock of Ir solution. This gives the Ir concentration in the HPLN/Ir solution in mg/mL. Loading efficiency can be determined by taking the Ir concentration in mg/ml divided by the HPLN concentration in mg/ml to yield mg Ir/ mg HPLN.

### *Anti*-CD19 or *anti*-CD99 conjugated micelle

Human monoclonal antibodies to CD19 or CD99 were partially reduced by treatment with TCEP. 1.0 mg of antibody is mixed in 1 mL of PBS with 5.0 mmoles TCEP and allowed to react at room temperature for 30 min on a low-speed sample mixer (HulaMixer, Invitrogen). 0.16 mg of mal-Peg2000-DSPE and 0.62 mg of m-Peg2000-DSPE are hydrated in the TCEP-reduced antibody solution. The reduced antibodies are reacted with lipid micelle for 4 hours at room temperature on the sample mixer. The reaction was quenched by adding 25 mmoles L-cysteine and incubating at room temperature for 30 minutes on the sample mixer. The reaction mix was centrifuged at 10,000 rpm for 10 minutes to pellet any insoluble debris. Removal of the TCEP and excess L-cysteine reagents was accomplished by spinning the solution in a 100 kDa MWCO centricon (Amicon Ultra-4 Ultracel, MilliporeSigma, Burlington, MA). The reagents were removed by doing four 10X PBS washes. The antibody concentration was measured by the 280 nm absorbance. Antibody concentration in the micelle in mg/mL=Abs_280_/1.4.

### NV102

1.78 mg of *anti*-CD19 labeled micelle was mixed with 35.64 mg HPLN/Dox and centrifuge at 10,000 rpm for 5 min. the supernatant is transferred to a fresh tube and allowed to rock on the sample mixer at ambient temperature for 8 hours. The solution is centrifuged at 10,000 rpm for 5 min. The HPLN and Dox concentrations are determined as described above. The NV102 antibody load is assessed by SDS-PAGE gel and ELISA, targeting the IgG component. The particle size is assessed on the Zetasizer Nano-ZS.

### NV101 and NV103

NV101 and NV103 are generated using *anti-*CD99 antibody in place of *anti-*C19 antibody, as described above. The HPLN and Dox or Ir concentrations are determined as previously described. Antibody load and particle size are assessed as previously described.

### Flow cytometry

To examine the binding efficiency of HPLNs, 1×10^6^ cells (REH cells or TC71 cells) were collected from confluent culture flasks by scrapping and incubated with 50*μ*g HPLNs in 1ml culture media for 1 hour at room temperature. The cells were vortexed in PBS and centrifuged at 2,000 rpm, repeating this process three times. The HPLN-labeled cells were then analyzed using BD LSR II flow cytometer (BD Biosciences) through a PECy5 filter using DiVA (version 4.1.2) software. Samples were first gated on intact cells by FSC and SSC profile. Mean fluorescence of PECy5 signal (geometric MFI, indicating surface-bound HPLNs) was measured in intact cells and was analyzed relative to that from untreated cells and cells treated with untargeted HPLNs as controls.

### Protein lysates and Western blotting

Ewing sarcoma cell line TC32 cells were seeded in 6-well plate and incubated overnight. The following day, cells were treated with 1uM of doxorubicin, HPLN/Dox or NV101 for a two-hour period then washed with fresh media. Cells were incubated for 24 hrs at 37^◦^C and harvested. Harvested cells were rinsed in PBS containing 100 mmol/L Na_3_VO_4_ and lysed in NP40 lysis buffer (50 mmol/L HEPES, 100 mmol/L NaF, 10 mmol/L Na_4_P_2_O_7_, 2 mmol/L Na_3_VO_4_, 2 mmol/L EDTA, 2 mmol/L NaMoO_4_, and 0.5% NP40) containing a Roche protease inhibitor cocktail for 30 min at 4°C with shaking. Protein concentrations were standardized using detergent compatible Bio-Rad protein assay kits. Standard Western blot analysis was done with antibodies to poly (ADP-ribose) polymerase.

### Caspase-3 assays

Caspase-3 activity was assayed by Z-DEVD-AFC cleavage according to the manufacturer’s protocols (Calbiochem). Briefly, cells grown under the indicated conditions were lysed at 4°C and lysates were incubated with reaction buffer containing 50 μmol/L Z-DEVD-AFC. After incubation at 37°C, fluorescence was measured with a GENiosPro instrument (Tecan). Caspase-3 activity was adjusted for protein concentration and relative activity was expressed as the fluorescence ratio between the normalized caspase-3 activities of nontreated cells (relative unit of 1.0) versus treated cells.

### Cell Viability Assay

REH or TC71 cells were grown in RPMI 1640 with 10% FBS and antibiotics (penicillin/streptomycin). Cells were seeded in a 96-well plate at a concentration of 1×10^4^ cells/well with a volume of 100 µL media and incubated overnight. The following day, cells were treated with NV101, NV102 or NV103, untargeted HPLN/dox, HPLN/Ir, Empty HPLNs, or free drugs. Doses were added based on free drug concentrations ranging on a log scale from 0.01 to 100 µM and at 0 nM. The 0 nM well was treated with PBS. Each treatment was performed in triplicate. Cells were incubated under standard CO_2_ conditions for 8, 24, and 72hrs at 37°C. The cell viability was measured using Cell Titer-Glo® Luminescent Cell Viability Assay Kit (Promega, USA) according to the manufacturer’s protocol.

### Xenograft model of primary leukemia/ NOD/SCID mouse leukemia or Ewing Sarcoma model

TC71 or primary ALL *LAX7R* cells were labeled with firefly luciferase (LUC) by transduction with pCCL-MNDU3-LUC viral supernatant and intravenously injected into sublethally irradiated (250 cGy) NOD/SCID mice (Jackson Laboratories) (0.05 x 10^6^cells/mouse). 8 days post tumor cell injection, five mice received only buffer, five received drug-filled no antibody HPLNs (**untargeted**) and five received NV101 (TC71 model) or NV102 (*LAX7R* model). The dosing schedule was conducted so that the animals received the same drug dose in HPLNs, two times per week at a dose of 2 mg/kg drug. Serial monitoring of tumor progression in mice was performed as described earlier [62] at indicated time points using an in vivo IVIS 100 bioluminescence/optical imaging system (Xenogen). D-Luciferin (Promega) dissolved in PBS was injected intraperitoneally at a dose of 2.5 mg per mouse 15 minutes before measuring the luminescence signal. General anesthesia was induced with 5% isoflurane and continued during the procedure with 2% isoflurane introduced via a nose cone. Moribund mice were sacrificed based on over 15% body weight loss of the animals. All mouse experiments were subject to institutional approval by Children’s Hospital Los Angeles IACUC (Institutional Animal Care and Use Committee). The log-rank test was used to evaluate differences in the median survival time (MST) for the in vivo studies. **P<0.005; ***P<0.0001 indicated a significant difference.

## Supplementary Data

### *In vitro* Ewing Sarcoma Treatment with NV101 and NV103

HPLNs that were Doxorubicin (Dox) loaded were mixed with *anti*-CD99 conjugated antibody micelles to produce the NV101. HPLNs that were Irinotecan (Ir) loaded were mixed with *anti*-CD99 conjugated antibody micelles to produce the NV103. TC71, a CD99-expressing Ewing sarcoma cell line, was chosen for the study. The binding of empty *anti*-CD99 antibody labeled HPLNs to TC71 cells was confirmed by FACS analysis **(see manuscript Figure 5**). Using the optimal Ab-micelle to HPLN ratio of 28 ug micelle to 1 mg HPLN/Dox, experiments were then performed to determine whether *anti-*CD99 targeting function improved the ability of Dox-loaded HPLNs to inhibit growth of Ewing sarcoma cells, *in vitro*. As a negative control, untargeted HPLN/Dox was prepared by inserting cysteine-terminated micelles into the nanoparticle membrane. When comparing NV101 and free doxorubicin at the 24-hour time point, we observed approximately 60% improvement in the targeted particle over free drug (**Supplementary Figure 1A**). At later time points the difference in potency between NV101 and untargeted HPLN/Dox or free drug became negligible. Empty HPLNs and empty CD99-targeted HPLNs did not demonstrate significant TC71 cell killing.

Similar to NV101, HPLNs that were Irinotecan (Ir) loaded were mixed with *anti*-CD99 conjugated antibody micelles to produce the NV103. Using the optimal Ab-micelle to HPLN ratio of 28 ug micelle to 1 mg HPLN/Ir, experiments were then performed to determine whether *anti-*CD99 targeting function improved the ability of Ir-loaded HPLNs to inhibit growth of Ewing sarcoma cells, *in vitro*. NV103 demonstrated a growth inhibitory potency over untargeted HPLN/Ir of approximately 53% at the 8-hour time point; 25% at the 24-hour time point and 44% at the 72-hour time point (**Supplementary Figure 1B**). In this *in vitro* assay, NV103 preformed demonstrably better in terms of cell cytotoxicity than either free irinotecan or untargeted HPLN/Ir at the two early time points (8 and 24 hours). At the latest time point tested, 72-hours, NV103 still slightly outperformed untargeted HPLN/Ir and became equivalent to free irinotecan. Again, as with the NV101 studies above, neither of the empty HPLNs or antibody-targeted, empty HPLNs demonstrated significant TC71 cell killing.

**Supplementary Figure 1.**
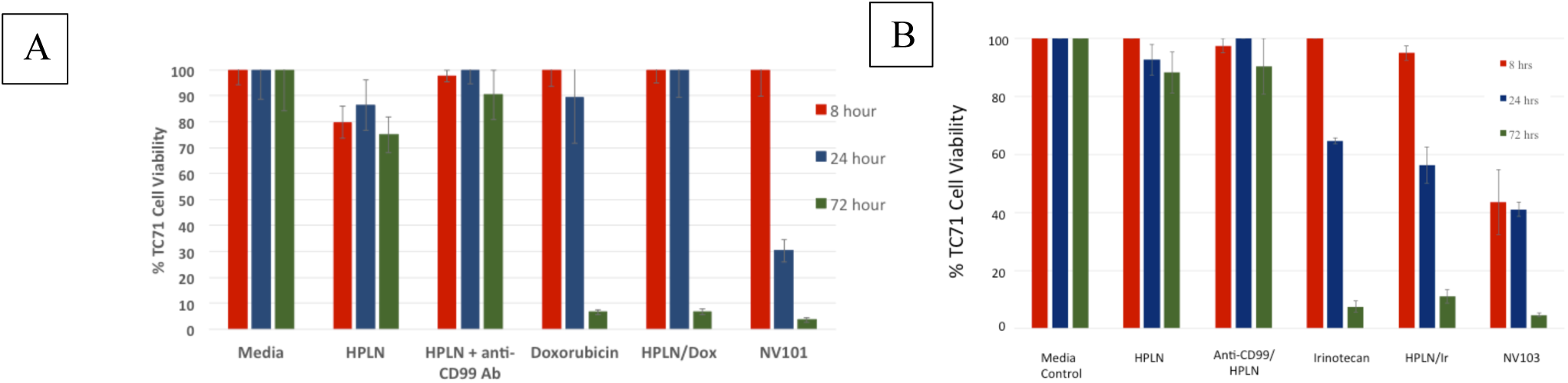
Human Ewing sarcoma TC71 cells were exposed to various untargeted, *anti-*CD99 targeted, empty nanoparticles and free drug formulations. The cytotoxicity was measured with an MTT assay. The 100% cell survival level was established as the media control percentage of live cells. All assays were done in triplicate. A: dox-loaded formulation: NV101; B: ir-loaded formulation: NV103.

### *In vitro* Leukemia Treatment with NV102

The doxorubicin loaded HPLNs, from which NV101 was prepared (above), were instead mixed with *anti*-CD19 conjugated antibody micelles to produce the NV102. REH, a CD19-expressing human leukemia cell line, was chosen for the study. The binding of empty *anti*-CD19 antibody labeled HPLNs to REH cells was confirmed by FACS analysis **(see manuscript Figure 5**). Using the optimal Ab-micelle to HPLN ratio of 28 ug micelle to 1 mg HPLN/Dox, experiments were then performed to determine whether *anti-*CD19 targeting function improved the ability of Dox-loaded HPLNs to inhibit growth of REH cells, *in vitro*. As a negative control, untargeted HPLN/Dox was prepared by inserting cysteine-terminated micelles into the nanoparticle membrane. NV102 demonstrated a growth inhibitory potency over untargeted HPLN/Dox of approximately 11% with REH cells at the 24-hour time point (**Supplementary Figure 2**). At later time points the difference in potency between NV102 and untargeted HPLN/Dox became negligible. With this cell line, in this this *in vitro* assay, free doxorubicin showed the most potency, consistent with previously reported models, where free doxorubicin was approximately 38- to 82-fold more potent than conventional liposomal doxorubicin [Supplementary reference 1]. The difference in the potency of the free drug verses liposomal drug is primarily due to delayed release of drug from the endocytic compartment of cells that have taken up liposomal doxorubicin [Supplementary reference 1]. Neither empty HPLNs or antibody-targeted, empty HPLNs had observable REH cell killing.

**Supplementary Figure 2.**
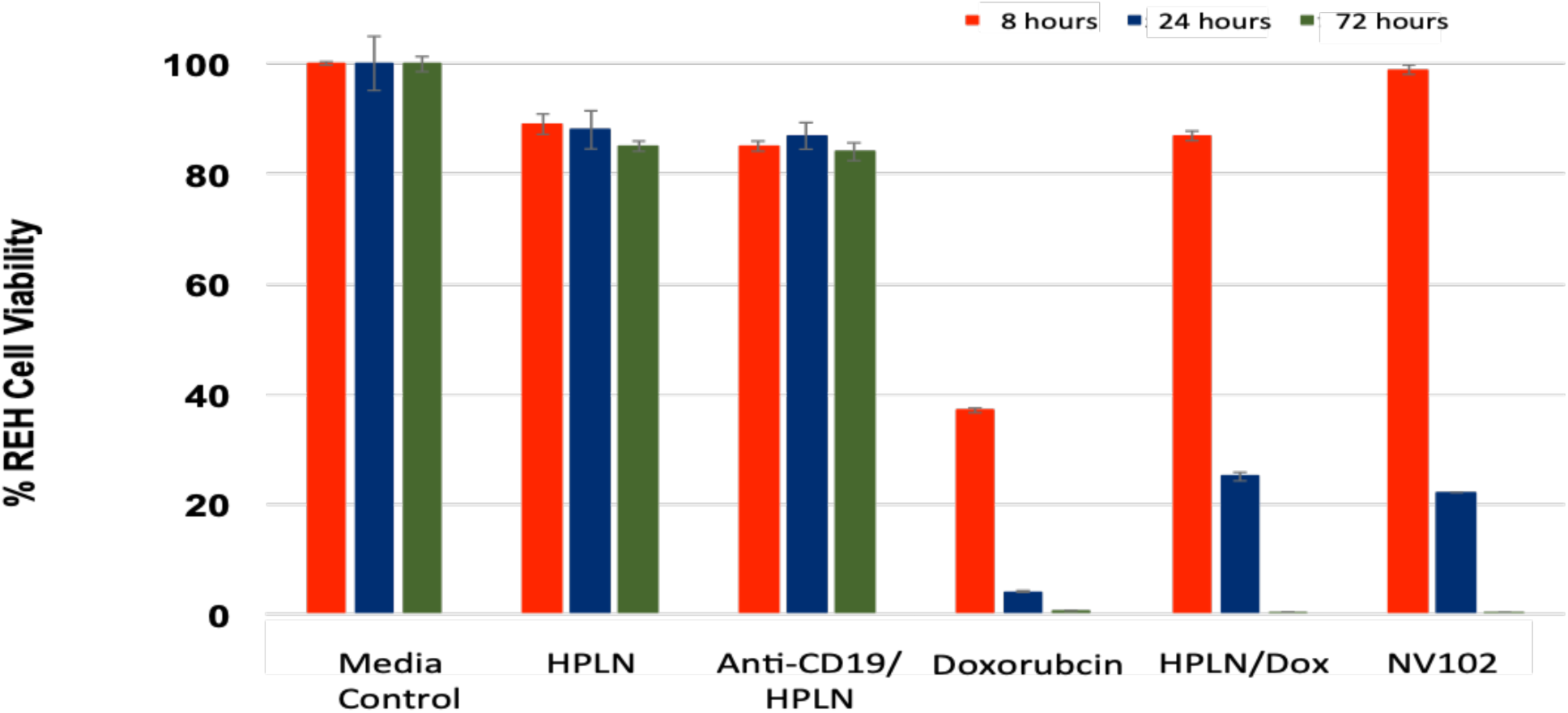
Human leukemia REH cells were exposed to various nanoparticle and drug formulations and the cytotoxicity was measured with an MTT assay. The 100% cell survival level was established as the media control percentage of live cells. All assays were done in triplicate.

### NV103 PK/PD studies

The toxicological and PK/PD parameters for NV103 were assessed in normal mice. For this study, free Ir injection was given at a 10-fold greater dose than the Ir contained in the NV103 particles. As can be seen in **Supplementary Figure 3 A**, the plasma concentrations of Ir vs. NV103 delivered Ir shows a much slower clearance of the drug encapsulated in the nanoparticle. This is seen for the observed Ir in the half-life, Cmax and AUC measurements **Supplementary Figure 3 B**. While the time to establish the maximal Ir concentration (Tmax) is similar for both NV103 and free Ir, the clearance of Ir described as the amount remaining in circulation (AUC) suggests that the circulation kinetics are greatly affected by the form administered (encapsulated vs. free), with the encapsulated form showing more than 5-fold increased AUC verses free Ir. Another analysis looked at the plasma levels of the active metabolite SN-38. SN-38 plasma levels for NV103 treated animals resulted in a greater Cmax and longer circulation half-life than in animals with free Ir. The final PK analysis assessed the Ir bone marrow concentration versus time profile. In contrast to the plasma Ir levels, the Ir derived from NV103 compartmentalized to bone marrow was **4-fold less** than free Ir. (Data not shown).

**Supplementary Figure 3.**
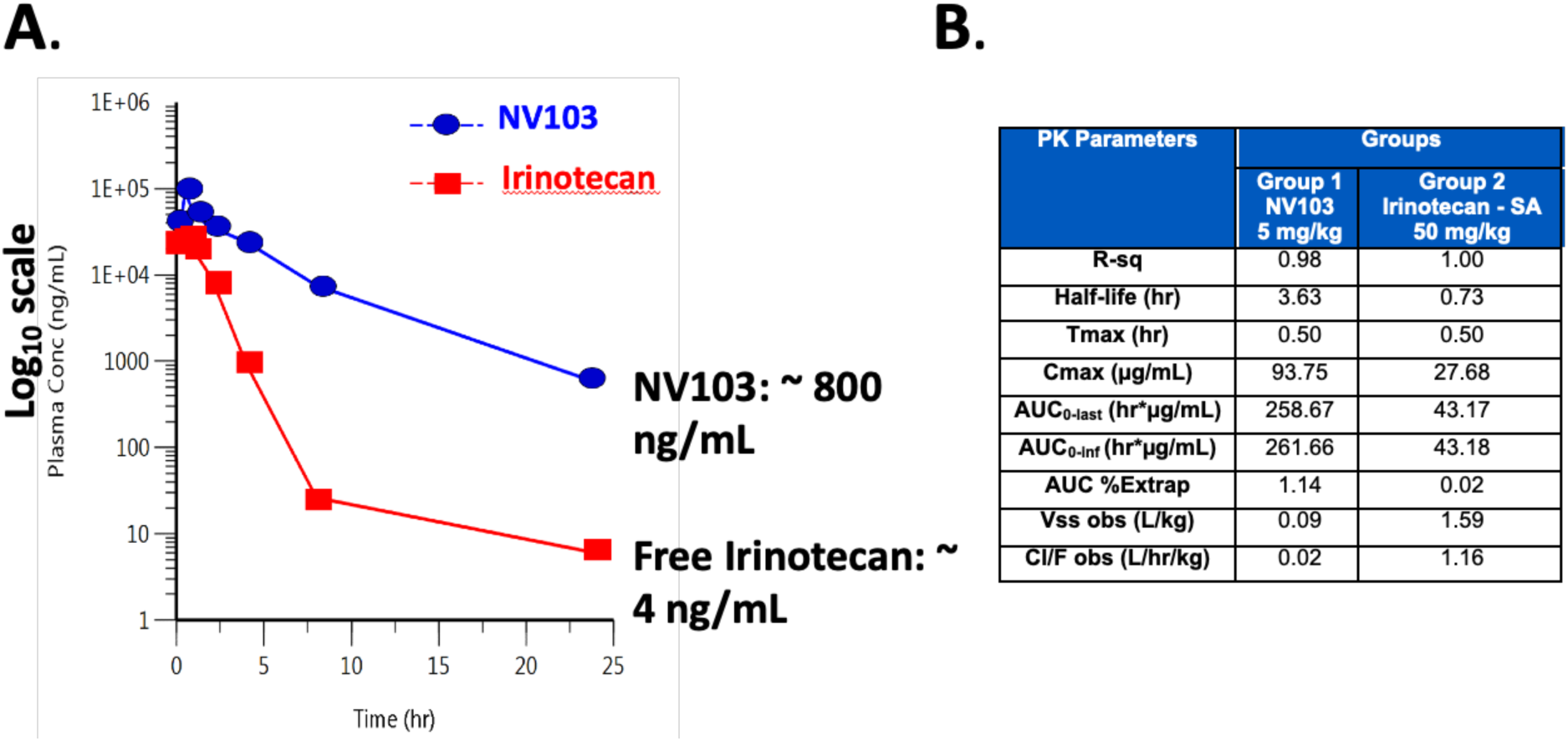
A: Normal mice plasma concentrations of irinotecan in mice treated by the free drug vs. irinotecan delivered in NV103.B: Observed PK parameters of NV103 (Group 1) vs. free drug (Group 2) in treated normal mice.

Key terms: Nanoparticle, cancer, polydiacetylene, Ewing Sarcoma, adult Leukemia, ALL, antibody targeting, Doxorubicin, Irinotecan, transgenic mice, CD99, CD19.

## Acknowledgements

The authors thank their colleagues Dr. Jodi Hedges and Ms. Hannah Certo for critical reading of the manuscript and personnel at Children’s Hospital Los Angeles for their technical support, discussions, and suggestions on this work.

## Author Contribution

All authors have seen and approved the version of the submitted manuscript.

## Brand names

Proprietary brand names of pharmaceutical products and prescription medications used in this article are owned by their respective trademark owners.

Article: © 2025 The Authors.

